# Walking the Tightrope: Balancing Opposing Cooperativities as an Operating Principle in Dynein Assembly

**DOI:** 10.1101/2025.07.14.664506

**Authors:** Douglas R. Walker, Lisa Otten, Mukhtar O. Idris, Brittany Lasher, Daniel M. Zuckerman, Elisar J. Barbar

## Abstract

The central region of the cytoplasmic dynein complex, comprising the intermediate chain (IC) and two light chains (LC8 and Tctex1), has eluded thorough quantitative characterization due to its participation in a highly coupled seven-state binding network. Although isothermal titration calorimetry (ITC) is the gold standard for measuring binding thermodynamics, conventional analyses are limited to simple interaction schemes because individual isotherms contain insufficient information to resolve complex reaction networks. Here, we overcome this limitation by combining extensive experimental sampling with hierarchical Bayesian inference. We collected 39 ITC isotherms spanning eight experiment types and developed a global Bayesian framework integrating multiple datasets while explicitly accounting for concentration uncertainty. Using this approach, we fit the complete dataset to a mechanistic seven-state model, estimating 190 parameters, including 12 thermodynamic parameters while marginalizing over 178 nuisance parameters. Remarkably, this strategy yields 95% confidence intervals for thermodynamic values as narrow as 0.05 kcal/mol and back-propagates to nanomolar precision in effective concentrations, even when experimental concentrations are in the hundreds of micromolar. The resulting thermodynamic landscape enables predictive modelling of assembly populations under different scenarios, including binding states inaccessible to standard ITC analyses. These results reveal previously unrecognized binding states that may play key roles in dynein cargo attachment and release. More broadly, this work reveals a form of “multi-cooperativity” governing dynein assembly and demonstrates how intensive experimentation coupled with modern statistical tools can resolve complex molecular systems beyond the reach of traditional biophysical techniques.

**Significance Statement:** Large, complex mechanistic processes have remained difficult to fully characterize, which limits interpretability of the underlying biology. We utilize a large dataset of 39 complementary experiments to fully characterize a seven-state system using Bayesian inference. This process achieves impressively precise fits with 0.05 kcal/mol width confidence intervals. The high precision enables assessment of simultaneous positive and negative cooperativity in the assembly of the dynein intermediate chain with its light-chain partners. Simulation of state populations suggests that this balancing cooperativity is finely tuned to allow access to a half-bound state which has been previously inaccessible quantitatively. Our approach is broadly applicable and supports an emerging principle of molecular regulation—negative cooperativity as a strategy for tuning responsiveness and dynamic control.

## Introduction

Multivalent protein interactions are widespread and play essential roles in biological regulation(1–4). While multivalency is often assumed to enhance function through positive cooperativity among binding sites, many systems instead exhibit neutral or even negative cooperativity, even when multiple binding events appear similar or contribute to the same functional outcome. A classic example is insulin binding to its receptor(5), where negative cooperativity ensures sensitivity across a wide range of insulin concentrations. Within the dynein motor complex, the interaction between the intermediate chain (IC, Fig. 1a) and its two dimeric light chain partners, LC8 and Tctex1(6–9), offers an ideal model for examining how positive and negative cooperativity can coexist and emerge from distinct molecular features(8–11). This system offers insights into how nature balances stability and flexibility in complex multivalent protein assemblies.

**Figure 1.**
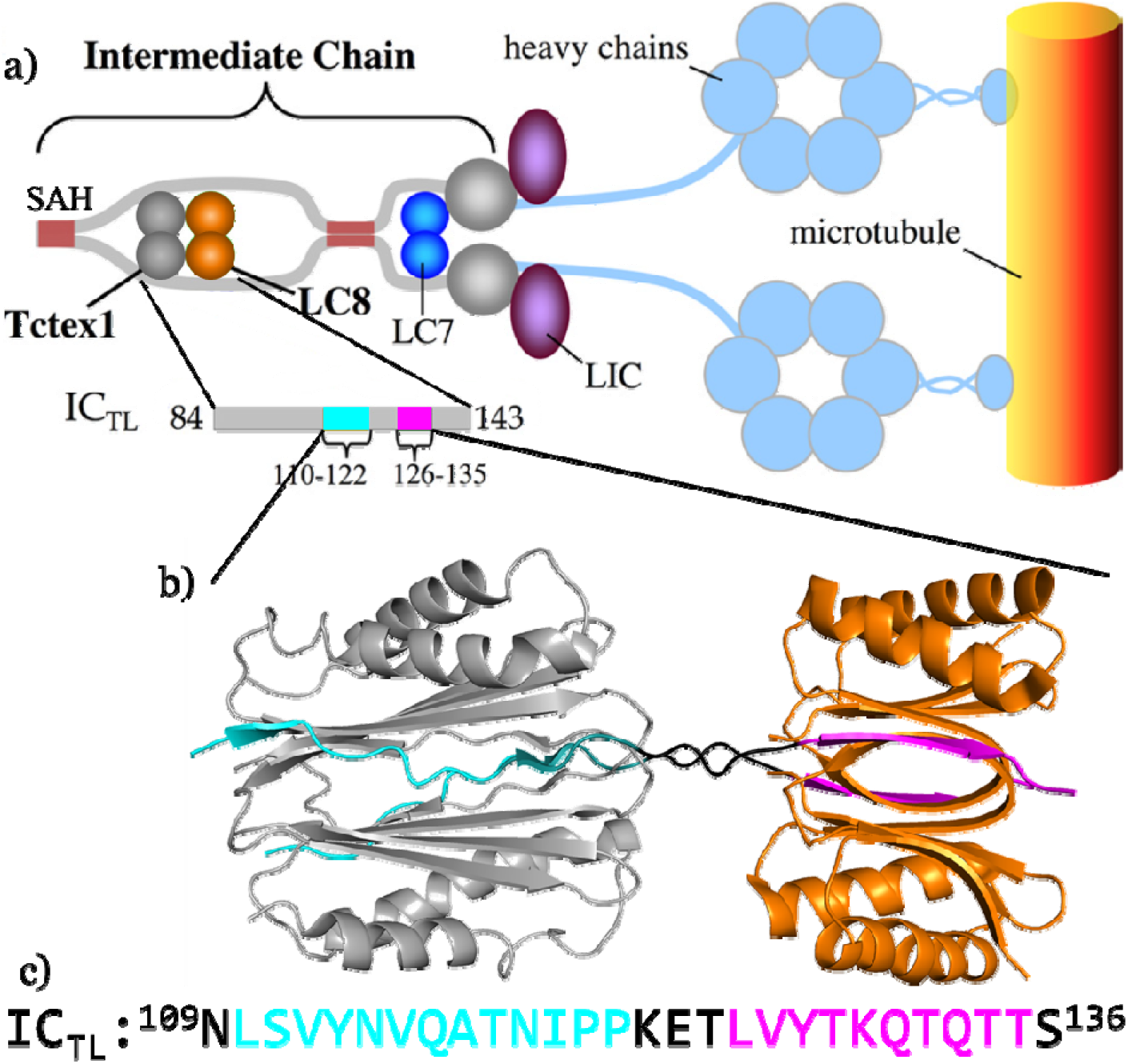
Overview of dynein intermediate chain IC with light chains LC8 and Tctex1. a) Cartoon of IC in the context of dynein to show the location of LC8 and Tctex1 binding, adapted from Hall et al.(8). b) The crystal structure (3FM7) of Tctex1 (grey) and LC8 (orange) bound to IC as reported by Hall et al.(8) illustrating a comparison of the two homodimers and the proximity of their binding sites on IC. c) The sequence of LC8 (magenta) and Tctex1 binding sites (cyan) highlighted matching the color scheme of panels a and b.

LC8 and its homolog Tctex1 both mediate dimerization of partner proteins through short recognition motifs embedded in intrinsically disordered regions. Each forms a rotationally symmetric (C_2_) intertwined homodimer and binds partners along grooves parallel to the axis of symmetry (Fig. 1b)(8). Within dynein IC, LC8 and Tctex1 binding sites occur in close proximity (Fig. 1c), and binding of one light chain enhances the affinity of the other(8, 9). LC8 is well-characterized(12), with over 100 known partners, many of which display positive cooperativity such that the second copy of the client binds with a higher affinity than the first(13). In contrast, Tctex1 remains less well understood, and its cooperative behavior has not been quantitatively defined. A notable structural distinction between the two light chains is their length—despite similar cross-sectional area, Tctex1 is approximately 40% longer than LC8 along the axis of symmetry, resulting in a correspondingly longer binding motif (Fig. 1b,c).

Previous studies established that LC8 and Tctex1 bind IC(7, 8, 14) and stabilize IC dimerization(9, 15), thereby contributing to assembly of the dynein complex. However, quantitative analysis of this system has been constrained by necessary model simplifications. To render the system tractable, prior work reduced the fully coupled binding network to a four-state model, which was further compartmentalized into four independent two-state equilibria (Fig. 2)(8). While this approach enabled approximate estimation of binding energetics and the cooperativity between LC8 and Tctex1 in binding IC, it necessarily obscured the behavior of the complete interaction network. Interestingly, replacing the native Tctex1 binding site in IC with an LC8 binding sequence (IC_TL_ mutated to IC_LL_) resulted in a dramatic increase in stability amounting to a 20-fold enhancement in affinity for the second homodimer compared to the wild type IC(8). This result suggests negative cooperativity present in the native system counteracts the positive effects of multivalency. Although provocative, such conclusions rest on simplified assumptions that preclude direct interrogation of the fully coupled, multistate landscape.

**Figure 2.**
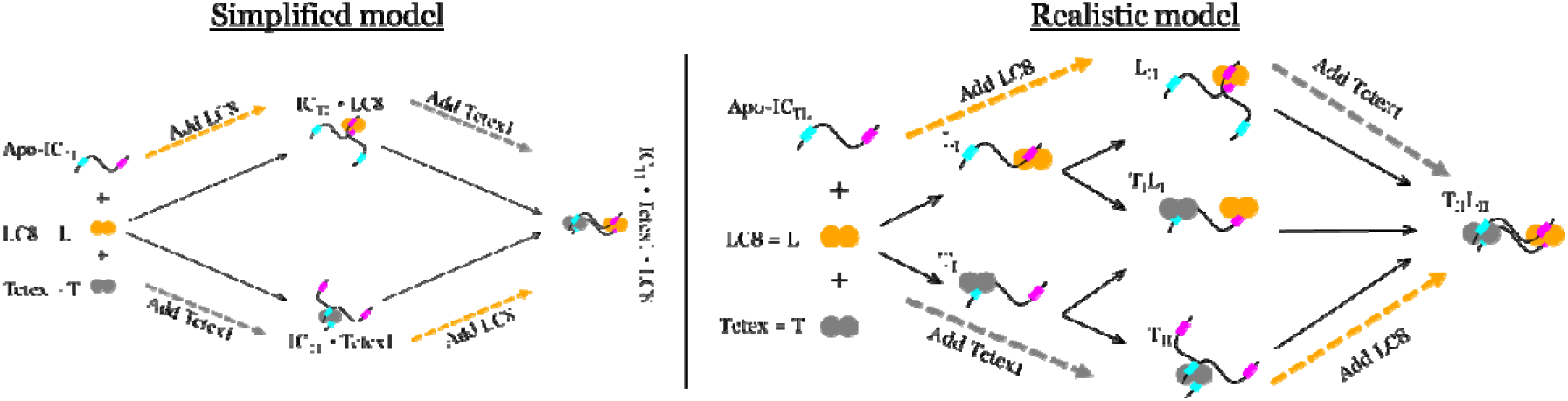
Simplified and seven-state models of the IC-LC8-Tctex1 network. (Left) Simplified four-state model used previously(8), which excludes several stoichiometric species. (Right) Realistic seven-state model of the IC/LC8/Tctex1 interaction. Dashed arrows schematically illustrate some of the experiments performed. Named states are referred to as labeled throughout the paper.

In this work we examine the IC-LC8-Tctex1 polybivalent system within a comprehensive thermodynamic framework. Rather than relying on the simplifying assumptions that constrained earlier studies, we adopt a full seven-state reaction model (Fig. 2) that more faithfully captures the molecular complexity and coupling inherent to this system.

Quantitative characterization of such a polybivalent network requires carefully designed experiments and rigorous data analysis. Measurements that probe reaction equilibria and binding energetics are inherently prone to error and uncertainty, particularly when limited information content and increasing system complexity impede robust data analysis. These challenges largely arise from “correlative effects” wherein multiple parameter sets produce effectively indistinguishable outputs(16–19). These challenges are amplified when the concentrations of the analytes cannot be measured with high precision, as is often the case in protein studies, because concentrations are often directly correlated with physical parameters of the model(20–22). In simple two-state reactions where A + B ↔ C, inaccuracy in analyte concentration can be identified and ameliorated: Because reaction stoichiometries are constrained to whole numbers, measured concentrations can often be adjusted to achieve precise, physically representative measurements for the reaction(20). However, introducing even a single additional state, as in a three-state model, removes the discrete constraints imposed by integer stoichiometries, allowing continuously variable ratios, enlarging the correlative effects associated with concentrations, in addition to those existing between physical parameters. Consequently, uncertainties in concentration measurements propagate through the model yielding broad thermodynamic uncertainty(13, 16, 23, 24). As reaction networks grow larger, the dimensionality of the parameter space expands dramatically, further increasing opportunities for correlated parameters and reducing mechanistic discriminability.

These limitations are particularly evident in Isothermal titration calorimetry (ITC) experiments. ITC analysis is simple when the system itself is simple(25, 26). For simple two-state systems, with a good quality isotherm and precisely measured concentrations, most thermodynamic parameters can be inferred almost by inspection, even without fitting software: enthalpy corresponds to the magnitude of the signal difference between early and late titration points, stoichiometry is indicated by the inflection point of the isotherm transition, and binding affinity is inferred from the slope of the line tangent to the inflection point(27). These few features are sufficient to fully characterize the system’s thermodynamics, and because stoichiometries are integer ratios, concentration-associated correlative effects can be minimal.

Incrementing the complexity of an ITC experiment to include a third state makes the relative contributions of each state to the overall shape of the isotherm unknowable via the simple heuristics noted above(25). A three-state system could result in a triphasic isotherm, or as more commonly observed, all three reactions could overlap resulting in a uniphasic isotherm(25, 28). Therefore, as the field increasingly studies multi-state systems, it becomes ever more important to have robust tools(13, 22, 24, 26, 29) for analyzing increasingly complex reactions and increasingly large datasets. For example, multivalent protein interactions often involve variable binding strengths and potential positive or negative cooperativity, which are challenging to parse, yet promise to provide a hotbed of interesting new scientific insight of which researchers have only just begun to explore(2, 30–34). To our knowledge, the largest model analyzed in the literature consists of four states(25) while the greatest number of isotherms globally analyzed is six(29).

Here, we overcome these limitations by applying hierarchical Bayesian inference (13, 24), to globally analyze 39 ITC isotherms spanning the full IC-LC8-Tctex1 binding network. This approach enables simultaneous estimation of all parameters while explicitly accounting for concentration uncertainties and mass-action constraints governed by Le Chatelier’s principle. Bayesian inference enables their uncertainties across the full parameter space, where each variable influences the others(22, 35, 36), an advantage that is particularly critical for ITC data where enthalpy-concentration correlations are strong and particularly insidious (13, 24, 25). By integrating multiple datasets, the hierarchical framework further improves precision and enforces consistency across experiments (24, 37).

Using this strategy, we estimate 190 parameters, including 178 nuisance and 12 thermodynamic variables, achieving 95% confidence intervals for thermodynamic values with widths as small as sub-percent. This exceptional precision enables several new insights. We show that IC is the least positively cooperative LC8 binding partner characterized to date, provide the first evidence that Tctex1 also exhibits a modest degree of positive cooperativity when dimerizing client proteins, and show that simultaneous binding of LC8 and Tctex1 to a single IC strand is likely negatively cooperative. Strikingly, the hierarchical Bayesian analysis also yields highly precise fits for the analyte concentrations, despite the size of the seven-state model, and the use of approximately 100 protein samples across the 39 isotherms. Leveraging these thermodynamic parameters, we further model the population distributions of all binding states under biologically relevant conditions. These simulations predict substantial populations of species in which LC8 and Tctex1 bind to only one IC strand, raising intriguing possibilities for their roles in dynein assembly and cargo transport. Together, these results highlight the intricate balance of positive and negative cooperativity in multivalent protein systems and demonstrate how Bayesian inference can reveal mechanistic insights inaccessible to conventional analysis.

## Results

### A More Realistic Model and 39 Isotherms

To capture the full complexity of the IC-LC8-Tctex1 interaction network, we expanded the previously used four-state model to a seven-state model (Fig. 2). The three additional species correspond to partially bound complexes (T_I_, L_I_, and T_I_L_I_) in which only one IC chain binds a light chain homodimer. These species contain a single IC molecule in the complex and a half-bound Tctex1 or LC8. Incorporating these “half-bound” states is essential because they arise naturally from mass-action equilibria and can be substantially populated depending on the experimental conditions and the underlying thermodynamics. We did not include intermediates corresponding to putative “open-ring” conformations of the fully bound T_II_L_II_ state in which either Tctex1 or LC8 is transiently half-bound. Such intramolecular interactions are concentration independent and therefore cannot be distinguished from the fully bound state by ITC. Their exclusion does not alter the equilibrium description of the system or the interpretation of the observable thermodynamic parameters.

A key advance of this work is that all relevant binding states are considered simultaneously in every experiment. Earlier analyses relied on simplifying assumptions that restricted each experiment to only a small subset of possible states, which made analysis tractable but did not reflect true equilibrium behavior. In reality, multiple species can coexist under most conditions, and their relative populations shift as experimental conditions change. Ignoring these additional states masks their contributions to the measured signal and limits accurate interpretation of cooperative effects within the interaction network

To characterize the energetics of IC binding to Tctex1 and LC8, we designed a targeted set of ITC experiments (Fig. 3) aimed at selectively enriching low-population intermediate species. These new measurements were combined with previously published data(8), summarized in Table 1, to enable comprehensive analysis of the full binding network. Earlier titrations placed IC in the calorimeter cell and were optimized to probe formation of the fully bound species T_II_, L_II_, and T_II_L_II_. To specifically enhance sensitivity of the partially bound intermediates T_I_, L_I_, and T_I_L_I_, we performed additional experiments with IC in the syringe. In total, our global analysis incorporated 39 ITC isotherms spanning eight experimental designs (Table 1 and Fig. 3), providing the diversity necessary to resolve all states in the seven-state binding system.

**Table 1.**
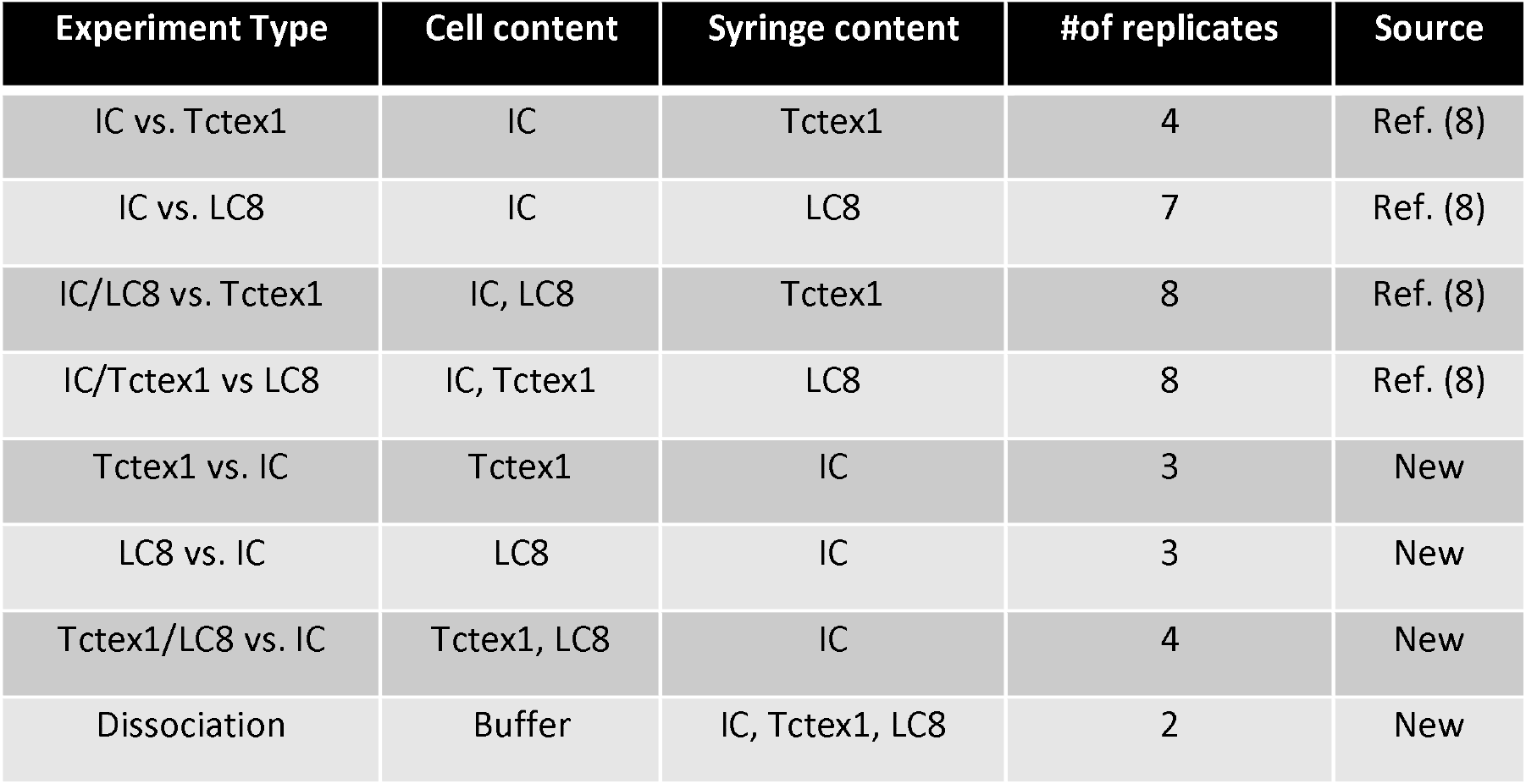
Inventory of experiments utilized for analysis of the IC-LC8-Tctex1 complex formation. For the “Experiment type” column, entries are formatted “cell” vs. “syringe”.

**Figure 3.**
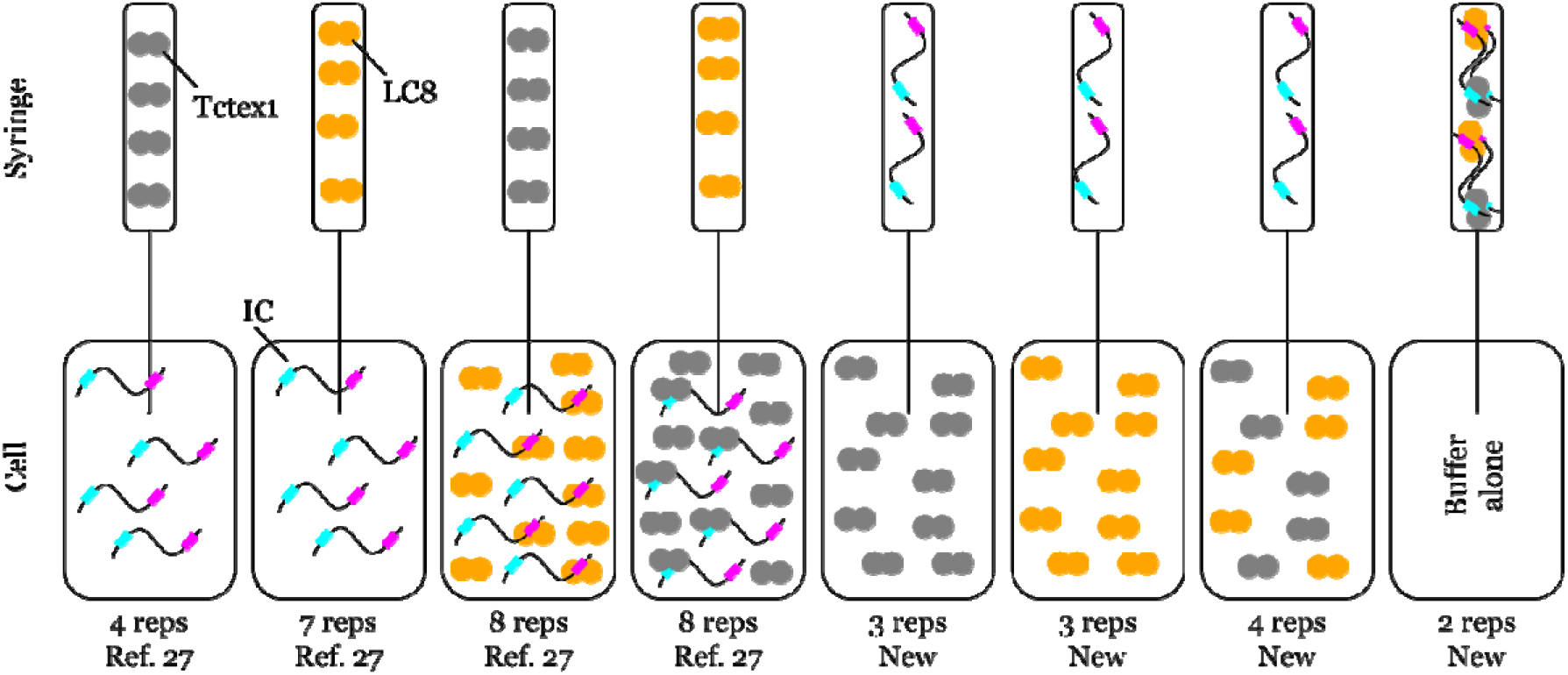
Schematic of the various ITC experiments that were collected and analyzed in the course of this work. IC, LC8, and Tctex1 are depicted visually as labelled.

### Targeted experiments and full-model fitting improve thermodynamic resolution

Figure 4 summarizes analysis of the simplest binding processes in the IC-Tctex-1-LC8 network (encircled in magenta), corresponding to bivalent binding of LC8 or Tctex1 to IC. Although these interactions appear conceptually straightforward, each forms a three-state system. As a result, the original 11 two-protein isotherms (Table 1 and Fig. 3)(8) were insufficient to fully resolve the energetics of bivalent binding. When the light chain dimers were titrated into IC, the doubly bound states (T_II_ and L_II_) were well constrained, but the singly bound intermediates (T_I_ and L_I_) remained poorly resolved.

**Figure 4.**
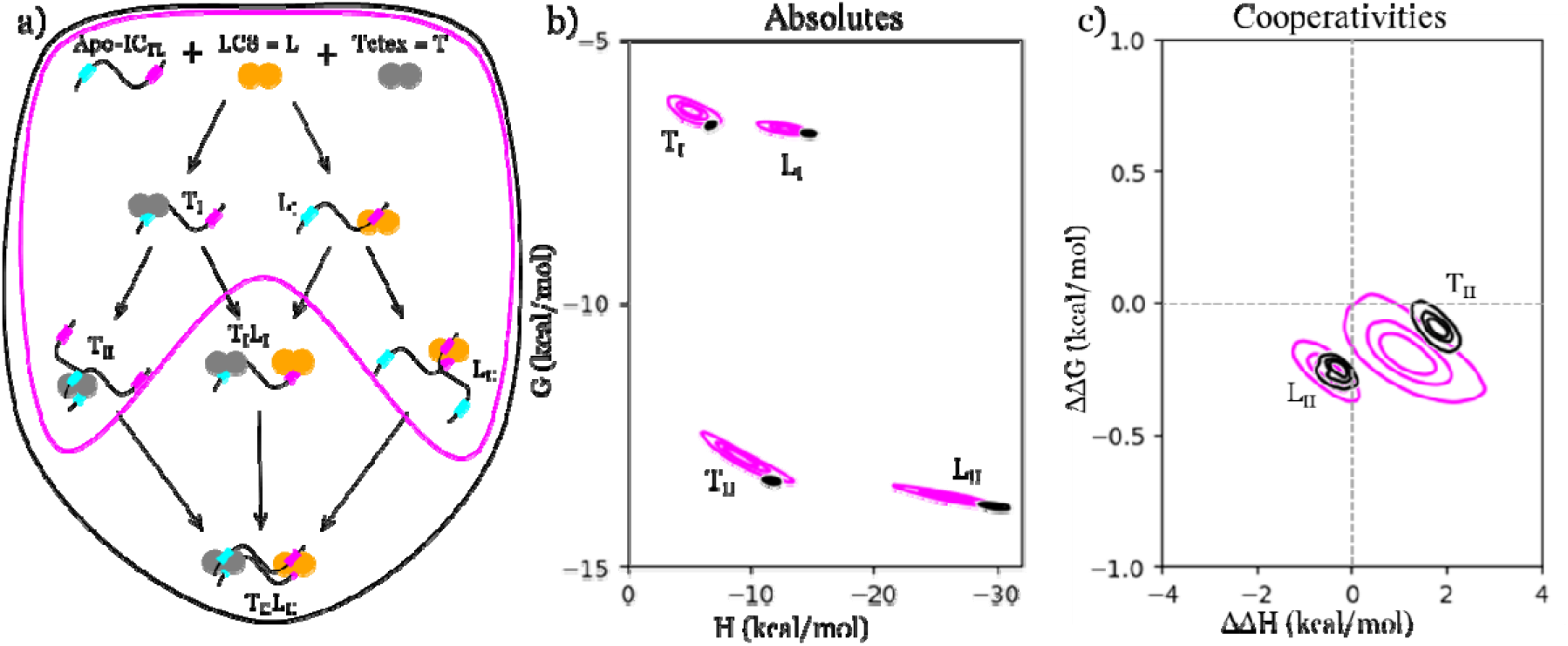
Thermodynamics of the simple (two-species) legs of the IC-Tctex1-LC8 binding network. Confidence contours from Bayesian inference are shown for fits using only the two-protein isotherms (magenta) and for global fits incorporating all isotherms (black). a) Seven-state model (rotated from Fig. 2) with the simple two-species legs highlighted in magenta. b) “Absolute” free energies and enthalpies for each interaction, referenced to the fully unbound state. c) Differences between the first (I) and second (II) binding events, defined as ΔΔG(T_II_) = (G(T_II_) - G(T_I_)) - G(T_I_). Dotted lines denote zero cooperativity. In all panels, contours enclose 39%, 68%, and 95% posterior probability.

To overcome this limitation, we collected additional “reverse direction” isotherms in which IC was titrated into the light chain homodimers, thereby increasing sensitivity of the singly bound states. This improvement arises from differences in species populations at early titration points. When LC8 is titrated into IC, IC is initially in excess, and both binding steps contribute to each injection. In contrast, titration of IC into LC8 places LC8 in excess, preferentially populating the half-bound intermediate and improving characterization of the first binding event.

We simultaneously fit all seven IC-Tctex1 isotherms and all ten IC-LC8 isotherms using our Bayesian pipeline. Thermodynamic parameters for the four species present in these subsets are shown in magenta contours in Fig. 4. Consistent with prior work(13), free energies were fit much more precisely than enthalpies. Inclusion of reverse-direction experiments significantly improved the free energy precision and modestly improved enthalpy estimates, underscoring the value of bidirectional titration strategies (13, 24). Thermodynamic parameters, confidence intervals, and cooperativity values derived from fitting the 17 two-protein isotherms are reported in Table S1.

Further refinement was achieved by globally fitting all 39 experiments in sequential batches as described in Methods. This full-model analysis markedly narrowed posterior distributions even for the simplest binding legs, as shown in black contours in Fig. 4 compared to the broader magenta contours derived from the two-protein subsets. This narrowing reemphasizes that accurate inference requires analyzing the complete equilibrium network, as all accessible species contribute to every experiment. The refined thermodynamics reveal IC as the least positively cooperative LC8 binding partner characterized to date and provide the first clear evidence for cooperativity in Tctex1 binding. Together, these results highlight the critical role of rigorous targeted experimental design combined with comprehensive Bayesian integration in resolving cooperative mechanisms in multivalent protein assemblies.

### Analysis of the full system reveals a mixture of positive and negative cooperativity

The final hierarchical Bayesian fits for the complete seven-state system are shown in Fig. 5. Incorporation of the three-protein isotherms (Table 1, Fig. 3) enabled resolution of the thermodynamics of the remaining states, T_I_L_I_, and T_II_L_II_, and substantially refined the thermodynamic parameters of the two-protein states (T_I_, L_I_, T_II_, and L_II_). Importantly, the central partially bound T_I_L_I_—unresolvable in earlier studies—was assigned thermodynamic values with high confidence, representing a key advance of this study.

**Figure 5.**
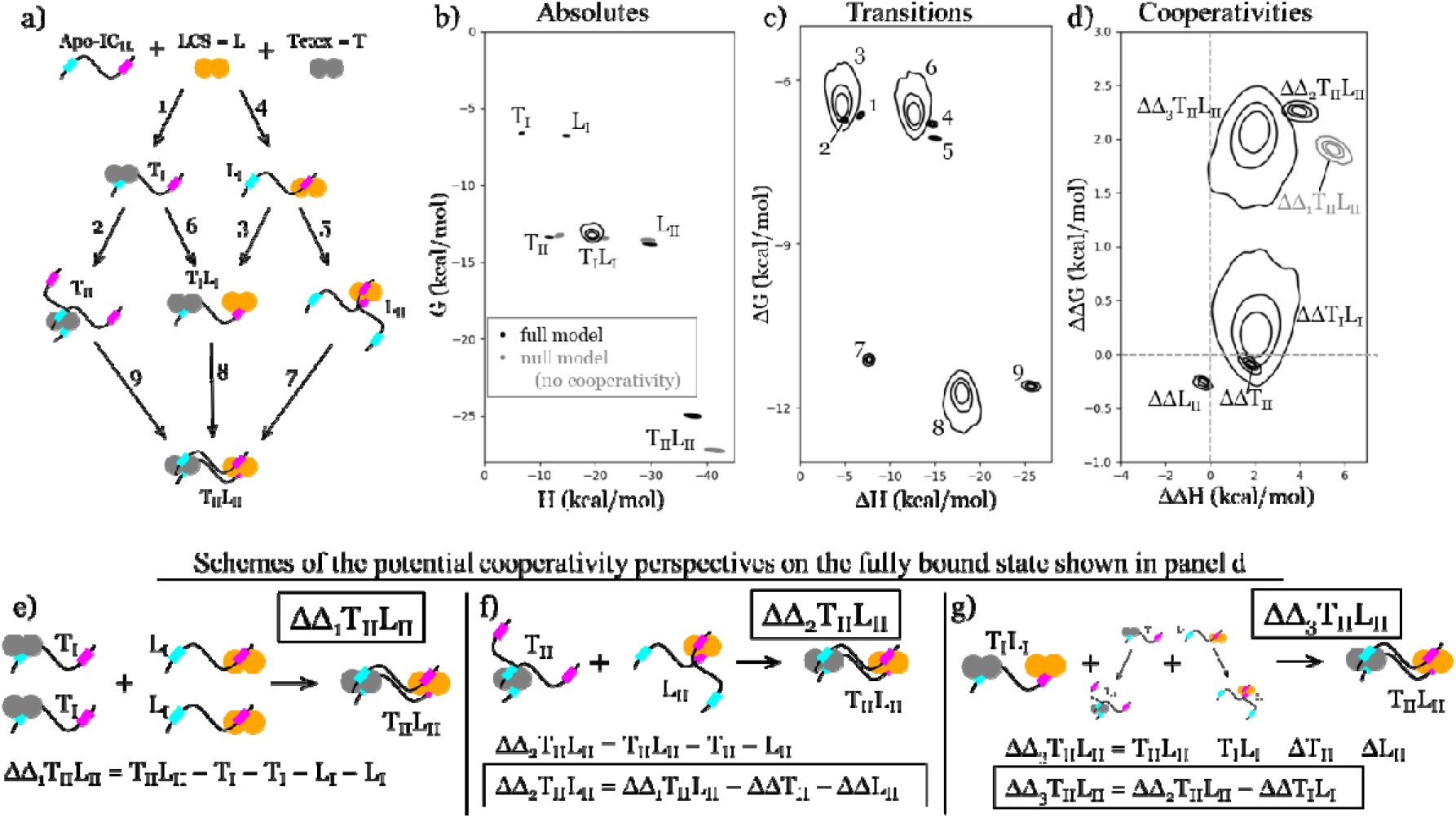
Thermodynamic analysis of the seven-state system reveals both positive and negative cooperativity. a) The seven-state model reoriented to align relative positions of each state with their corresponding energetics shown in panel b. b) Absolute enthalpy versus free energy for each state, referenced to the fully unbound state. Cooperativity is illustrated by comparing the Bayesian posterior derived from all data (black contours) with a non-cooperative null-model (gray contours). c) Free energy and enthalpy changes associated with each transition numbered in panel a, calculated as differences between the absolute values of the initial and final states connected by each arrow. d) Cooperativities in each state, defined as differences between the measured energetics and those expected from the constituent parts, as in Figure 4. For the fully bound T_II_L_II_ complex, three alternative decompositions of the constituent parts are shown, corresponding to the schematics in panels (e-g). Dotted lines indicate cooperativity=0. Values previously shown in Fig. 4 (black) are replotted for comparison. In all panels, contours depict 39%, 68%, and 95% enclosed posterior probability from the Bayesian analysis.

Overall, the thermodynamic parameters are consistent with previous measurements(8), but exhibit markedly improved precision and provide insight into three additional binding states. Thermodynamic parameters and cooperativities for all transitions as shown in Fig. 5c and d, with corresponding 95% confidence intervals listed in Table 2. Absolute free energies and enthalpies (Fig. 5b), including null model values, are reported in Table S2. Among the three multi-component states (T_II_, T_I_L_I_, and L_II_), free energy analysis shows that both dimerized states exhibit modest positive cooperativity, whereas T_I_L_I_ trends toward negative cooperativity. In contrast, the enthalpic contributions differ substantially across states: only L_II_ shows favorable enthalpic cooperativity, while both T_II_ and T_I_L_I_ exhibit unfavorable enthalpic contributions. Although LC8 binding is entropically disfavored overall, entropy does not contribute to LC8 cooperativity. However, both T_II_ and T_I_L_I_ transitions are entropically cooperative, and this favorable entropic contribution drives the net cooperativity observed for Tctex1 binding, largely compensating for the unfavorable enthalpy in the T_I_L_I_ transition (Table S3).

**Table 2.**
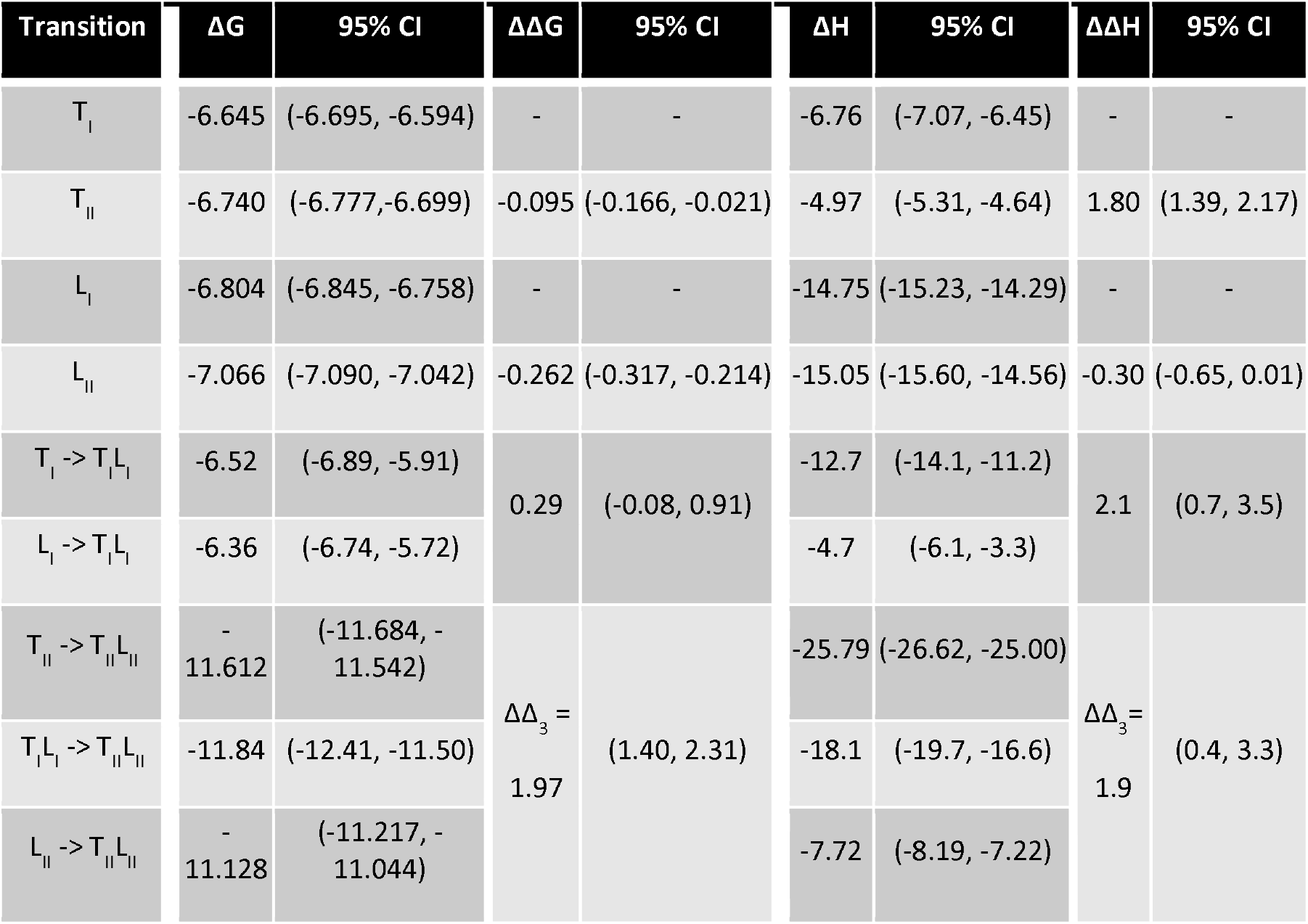
Measured thermodynamic values (in kcal/mol) and 95% confidence interval ranges for all transitions and cooperativities based on all isotherms.

For the fully bound T_II_L_II_ complex, comparison of the expected free energy and enthalpy if there was no cooperativity (Fig. 5b in gray) with the observed values (Fig. 5b in black) reveals substantial negative cooperativity in this system as explicitly shown in Fig. 5d. Interestingly, the ΔΔG value for the fully bound state (Table 2) closely matches the 2 kcal/mol difference in binding reported in Hall *et al*.(8) between IC_TL_ and IC_LL_ complexes. This correspondence suggests that the negative cooperativity quantified here may fully account for the observed differences in stability between these dynein subcomplexes.

Cooperativity defined relative to different reference states provides complementary insights into the forces governing assembly of the fully bound T_II_L_II_ complex. The simplest measure, schematized in Fig. 5e, compares the observed thermodynamics of T_II_L_II_ to those expected if each of the four binding events contributed independently. This metric, shown in grey in Fig. 5b,d, clearly indicates negative cooperativity. However, it double counts the positive cooperativity intrinsic to the formation of both the T_II_ and L_II_ subcomplexes and therefore underestimates the magnitude of negative cooperativity in the fully assembled state.

Additional insight is gained from employing other reference states that systematically remove these contributions. As illustrated in Figure 5f, the quantity ΔΔ_2_ represents the additional energetic cost of binding a second IC while discounting the stabilizing effects of binding each site on LC8 and Tctex1. The ΔΔ_3_ metric (Fig. 5d,g and Table 2) further removes the cooperative contribution arising from simultaneous binding of LC8 and Tctex1 to a single IC chain. While these corrections primarily broaden the free energy distributions, their impact on enthalpy is more pronounced, shifting the center of the ΔΔ_3_ distribution toward values consistent with little or no entropic contribution to cooperativity.

We also evaluated additional reference schemes, ΔΔ_4_ and ΔΔ_5_ (Fig. S1), which remove other sources of cooperativity. Across all five perspectives, the fully bound T_II_L_II_ complex consistently exhibits negative cooperativity in free-energy (Table S4), underscoring the robustness of this conclusion to the choice of reference state.

The full Bayesian analysis also estimates a large number of “nuisance” parameters that nevertheless carry physical meaning, including analyte concentrations, many of which were resolved with remarkably high precisions. Concentration posteriors yielded 95% confidence intervals as narrow as 230 nM. As a percentage of the fit concentrations, the tightest 95% confidence interval gave a range of +/-3% while the geometric mean of 95% confidence intervals is +/-10% of the measured concentration. Representative experimental isotherms, final model fits, and concentration posterior corner plots are provided in the SI.

### Modelling state populations reveals physiological accessibility of the central T_I_L_I_ intermediate

The high precision of the free energy estimates enabled calculation of the populations of all seven binding states at arbitrary total concentrations of IC, LC8, and Tctex1, parameters that directly influence physiological behavior. We simulated equilibrium populations using 10 µM IC, representative of the lower end of our ITC concentration range, across a broad spectrum of LC8 and Tctex1 concentrations. Figure 6 presents results computed using the full thermodynamic model (Fig. 6 panels a, b) alongside calculations in which the negative cooperativity term (ΔΔ_2_) for T_II_L_II_ is omitted for comparison (Fig. 6c and d). The concentrations and associated uncertainties plotted in panels a and c were selected based on the LC8 and Tctex1 concentrations at which each of the six states of interest reaches maximal populations in panels b and d.

**Figure 6.**
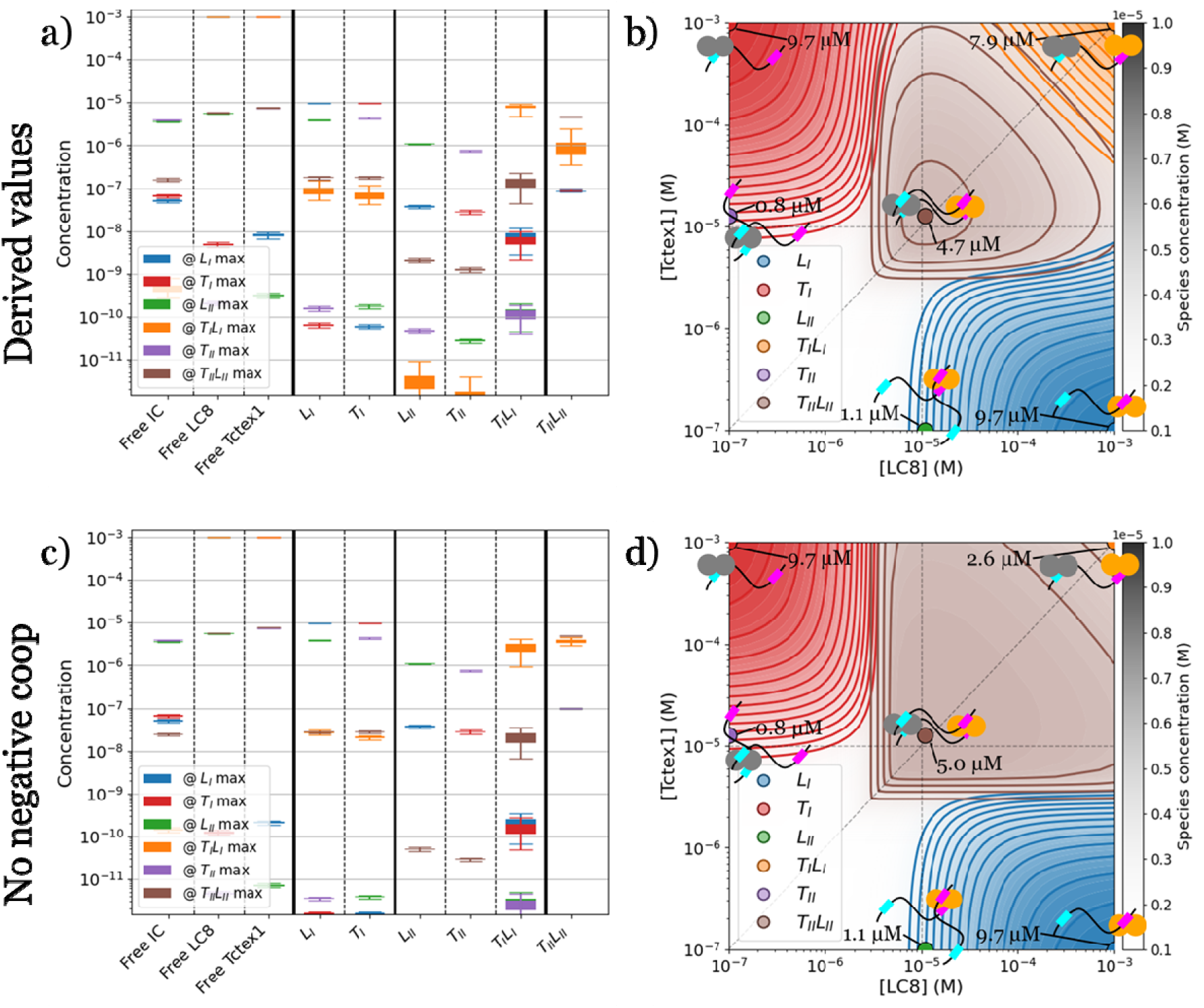
Effects of cooperativity on state populations. State-specific concentrations as functions of total concentrations of LC8 and Tctex1 are compared with a system lacking cooperativity. a) Concentrations and associated uncertainties for seven selected combinations of LC8 and Tctex1, calculated using Bayesian-inferred thermodynamic parameters. Each point corresponds to the condition that maximizes the population of one of the seven states over the matrix of LC8 and Tctex1 concentrations examined in panel b. b) Contour plots showing equilibrium concentrations of each of the seven states as [LC8] and [Tctex1] vary from 100 nM to 1 mM. c,d) Same analyses as in panels a and b, respectively, but for a non-cooperative model in which the free energy of the fully bound state is adjusted to remove cooperativity adjusted to remove cooperativity, G_null_(T_II_L_II_) = G(T_II_) + G(L_II_). In all panels, [IC] = 10 *µ*M. Dotted lines in panels b and d denote pairwise equimolar concentrations of LC8 and Tctex1.

These simulations provide both internal consistency checks and new mechanistic insights. In both models, the fully bound T_II_L_II_ complex reaches maximal population near equimolar concentrations of all three proteins (Fig. 6a). The doubly bound states, T_II_ and L_II_, peak at about 20% of the total IC concentration (Fig. 6b), reflecting a strong thermodynamic drive, via Le Chatelier’s principle, toward accumulation of the half-bound intermediates, T_I_ and L_I_. Strikingly, at sufficiently high concentrations of LC8 and Tctex1, T_II_L_II_ is no longer the major species. Instead, the central doubly half-bound state T_I_L_I_ becomes highly populated, reaching ~ 80% of the total IC concentration when LC8 and Tctex1 are present at 100-fold (Fig. 6b). At approximately 11-fold excess, T_I_L_I_ and T_II_L_II_ contribute roughly equally to the total IC population. As a computational control, removal of negative cooperativity reverses this trend. In this scenario, T_II_L_II_ remains the major species even at 100-fold excess of LC8 and Tctex1 (Fig. 6c), while T_I_L_I_ accounts for only ~25% of the total IC population (Fig. 6d), demonstrating that negative cooperativity is essential for stabilizing the T_I_L_I_ intermediate.

Importantly, T_I_L_I_ may be even more physiologically accessible than these simulations suggest. Dynein operates within a heterogeneous cellular environment in which local concentrations of IC and LC8 vary widely across different cellular compartments, along microtubules, and during different stages of the cell cycle(38–40). Molecular crowding is also expected to increase effective concentrations and favor partially bound states (41, 42). Simulations performed using reported intracellular concentrations of each species (43) yield similar results (Fig. S2), reinforcing the conclusions that T_I_L_I_ can reach substantial occupancy under biologically relevant conditions.

## Discussion

Dimerization of client proteins by the dynein light chains LC8 and Tctex1 is a conserved and widespread regulatory mechanism(6–8), yet quantitative characterization of the full ensemble of states accessed during multivalent assembly has remained challenging. Recent investigations have established the importance of distinguishing half-bound from fully bound LC8 complexes for characterizing the cooperativity across the LC8 dimer(13). Here, we extend this mechanistic resolution to the complete polybivalent IC-LC8-Tctex1 system by combining targeted ITC experiments with hierarchical Bayesian Inference to globally fit 39 isotherms to a biologically realistic seven-state model. This approach required fitting twelve thermodynamic parameters, 78 error parameters, and 100 protein concentrations, for a total of 190 parameters. Despite the size and complexity of the network, this approach yielded unusually precise thermodynamic parameters. Free energies of six of the nine transitions were resolved to 95% confidence interval half-widths below 0.1 kcal/mol, while similarly six of the nine enthalpy transitions were resolved to 95 % confidence interval half-widths less than 1 kcal/mol. This precision enabled direct interrogation of cooperative interactions that were previously inaccessible.

The resulting thermodynamic landscape reveals a mixture of positive and negative cooperativity. Assembly of the fully bound T_II_L_II_ complex slightly favors LC8 binding before Tctex1, though both pathways are energetically accessible. Importantly, the central T_I_L_I_ intermediate— previously unresolvable—emerges as a key state that becomes substantially populated at elevated LC8 and Tctex1 concentrations. Population modeling shows that negative cooperativity between LC8 and Tctex1 is essential for stabilizing T_I_L_I_; removing this effect abolishes its accumulation. These results suggest that negative cooperativity is not a design flaw but a functional feature, potentially enabling IC to act as a concentration-sensitive switch that regulates cargo release at the N-terminal single α-helix.

The broader ensemble-of-states behavior reinforces the functional relevance of half-bound intermediates in multivalent systems. Across much of the concentration space, singly bound states (TI and LI) dominate over fully dimerized complexes (TII and LII), implying that half-bound species may represent primary functional states rather than transient intermediates in LC8- or Tctex1-containing assemblies. Explicit consideration of such intermediate species will be essential for interpreting cooperativity and regulatory mechanisms in other multivalent networks. Beyond dynein biology, this study demonstrates the power of hierarchical Bayesian inference for extracting high-precision thermodynamic insights from complex, low-information systems. The approach(24) is broadly applicable to other techniques—not limited to ITC—provided that diverse experimental conditions can be sampled. By globally integrating diverse experiments and rigorously propagating uncertainty across all parameters, the Bayesian framework offers a robust solution for analyzing multivalent interactions, cooperative binding, and other intricate biochemical networks. As large, modular, and intrinsically disordered protein complexes continue to dominate modern cell-biological research, such approaches will be indispensable for revealing mechanistic principles governing their organization and regulation.

The intensive Bayesian computations and experimental design merit discussion. First, this study benefitted from an iterated design cycle in which initial Bayesian calculations, starting from existing isotherms(8) guided the selection of new experiments until sufficient precision was achieved to interpret the full seven-state network. Second, the initial Bayesian analyses were also used to generate synthetic isotherms at different conditions, enabling identification of the most informative experiments for improving parameter resolution. Third, the computational demands of fitting such a large dataset were substantial, requiring careful strategy to achieve convergence even using state-of-the-art parallelized *pocoMC* sampling (44). As detailed in the Supplementary Information, we employed two complementary strategies(45): (i) fitting subsets of isotherms to generate informative priors for subsequent analyses; and (ii) dividing the remaining isotherms into balanced sets and combining posterior samples through an annealing-based approach to approximate a global posterior. Together, these strategies enabled comprehensive inference across a model complexity well beyond that accessible to conventional approaches.

## Conclusions

By applying a recently refined and validated hierarchical Bayesian framework(24) to 39 ITC datasets, we achieved high-precision thermodynamic characterization of a seven-state binding network integral to the dynein light chain-dependent assembly. The analysis yielded multiple mechanistic insights into how LC8 and Tctex1 jointly shape a complex landscape of positive and negative cooperativity, including the previously overlooked functional importance of half-bound intermediates. We further identified the least positively cooperative LC8 client characterized to date and for the first time, provide a quantitative measure of cooperativity for Tctex1 binding. Notably, we resolve two negatively cooperative transitions within the same complex illustrating the intricate balance of stabilizing and destabilizing interactions that nature has encoded in its design of the dynein intermediate chain.

Our results suggest a plausible mechanism for dynein cargo release. By tempering the stabilizing effects of multivalency, combined LC8 and Tctex1 binding enables partial dissociation of IC dimers under conditions of limiting IC concentration. Such partial dissociation could facilitate cargo unloading at specific intracellular locations. More broadly, this behavior exemplifies an emerging principle in molecular regulation: negative cooperativity is not an evolutionary flaw, but a regulatory strategy widely used to fine-tune protein function, often in systems where precise and adaptable control is essential(5, 46–48).

In summary, this work clarifies the energetics underlying IC–LC8–Tctex1 assembly, uncovers a biologically plausible mechanism for dynein cargo release, and establishes a generalizable strategy for investigating multivalent interactions and cooperativity in complex biological networks. We anticipate that this framework will enable discovery of additional systems in which nature exploits the balance between positive and negative cooperativity to regulate molecular behavior with extraordinary precision.

## Materials and Methods

### Protein expression, purification, and preparation

*D. melanogaster* IC_TL_, LC8, and Tctex1 were prepared as described previously(9, 14, 49, 50). Purity was verified by SDS-PAGE and Size Exclusion Chromatography. Protein concentrations were calculated using a NanoDrop absorbance at 280 nm.

### Isothermal Titration Calorimetry

Experiments were performed in buffer composed of 50 mM sodium phosphate, 50 mM sodium chloride, and 1mM sodium azide at pH of 7.5. Proteins were co-dialyzed overnight prior to experiments to minimize heats of dilution occurring due to even slight buffer mismatch. A VP-ITC calorimeter (MicroCal, Northampton, MA) was used to measure binding thermodynamics at 25 °C. The design of legacy experiment syringe and cell components and concentrations were reported previously(8), however, more old experiments were used herein than were reported then. Further, we have found that intentional variation of conditions allows for pulling more information out of the same number of datasets(24, 51). Therefore, Table 3 lists the conditions of each of the experiments gathered and used in this work.

**Table 3.** List of experiments and conditions of ITC analyzed. T = Tctex1, L = LC8, I = IC. Concentrations of L and T are expressed as dimer concentrations. *Indicates where measured concentration was not previously reported explicitly, but just as being 5 times excess(8).

| Experiment | Syringe ($\mu\text{M}$) | Cell ($\mu\text{M}$) | Injection strategy | Source | BI Group |
| --- | --- | --- | --- | --- | --- |
| IC v. Tctex1 | 400 T | 80 I | 2 $\mu\text{L}$ + 27 (10 $\mu\text{L}$) | Ref. 27 | T prior |
| IC v. Tctex1 | 400 T | 80 I | 2 $\mu\text{L}$ + 27 (10 $\mu\text{L}$) | Ref. 27 | T prior |
| IC v. Tctex1 | 340 T | 60 I | 3 $\mu\text{L}$ + 27(10 $\mu\text{L}$) | Ref. 27 | T prior |
| IC v. Tctex1 | 400 T | 70 I | 3 $\mu\text{L}$ + 27(10 $\mu\text{L}$) + 7 $\mu\text{L}$ | Ref. 27 | T prior |
| IC v. LC8 | 250 L | 30 I | 2 $\mu\text{L}$ + 27(10 $\mu\text{L}$) | Ref. 27 | L prior |
| IC v. LC8 | 250 L | 50 I | 2 $\mu\text{L}$ + 27(10 $\mu\text{L}$) | Ref. 27 | L prior |
| IC v. LC8 | 400 L | 80 I | 2 $\mu\text{L}$ + 27(10 $\mu\text{L}$) | Ref. 27 | L prior |
| IC v. LC8 | 400 L | 80 I | 2 $\mu\text{L}$ + 27(10 $\mu\text{L}$) | Ref. 27 | L prior |
| IC v. LC8 | 375 L | 65 I | 3 $\mu\text{L}$ + 26(10 $\mu\text{L}$) | Ref. 27 | L prior |
| IC v. LC8 | 410 L | 54.6 I | 3 $\mu\text{L}$ + 29(7 $\mu\text{L}$) | Ref. 27 | L prior |
| IC v. LC8 | 400 L | 80 I | 3 $\mu\text{L}$ + 27(10 $\mu\text{L}$) + 7 $\mu\text{L}$ | Ref. 27 | L prior |
| IC/LC8 v. Tctex1 | 400 T | 80 I + 200 L* | 2 $\mu\text{L}$ + 27(10 $\mu\text{L}$) | Ref. 27 | Full 2 |
| IC/LC8 v. Tctex1 | 200 T | 80 I + 87.5 L* | 5 $\mu\text{L}$ + 27(10 $\mu\text{L}$) | Ref. 27 | Full 1 |
| IC/LC8 v. Tctex1 | 100 T | 20 I + 200 L* | 5 $\mu\text{L}$ + 20(10 $\mu\text{L}$) | Ref. 27 | Full 2 |
| IC/LC8 v. Tctex1 | 10 T | 2 I + 5 L* | 5 $\mu\text{L}$ + 23(10 $\mu\text{L}$) | Ref. 27 | Full 1 |
| IC/LC8 v. Tctex1 | 223.5 T | 15 I + 37.5 L* | 3 $\mu\text{L}$ + 22(10 $\mu\text{L}$) | Ref. 27 | Full 1 |
| IC/LC8 v. Tctex1 | 150 T | 23 I + 57.5 L* | 3 $\mu\text{L}$ + 24(8 $\mu\text{L}$) | Ref. 27 | Full 2 |
| IC/LC8 v. Tctex1 | 150 T | 23 I + 57.5 L* | 2 $\mu\text{L}$ + 39(5 $\mu\text{L}$) | Ref. 27 | Full 1 |
| IC/LC8 v. Tctex1 | 190 T | 20 I + 50 L* | 3 $\mu\text{L}$ + 29(5 $\mu\text{L}$) | Ref. 27 | Full 2 |
| IC/ Tctex1 v. LC8 | 400 L | 80 I + 200 T* | 2 $\mu\text{L}$ + 14(10 $\mu\text{L}$) | Ref. 27 | Full 2 |
| IC/ Tctex1 v. LC8 | 400 L | 80 I + 200 T* | 2 $\mu\text{L}$ + 16(10 $\mu\text{L}$) | Ref. 27 | Full 1 |
| IC/ Tctex1 v. LC8 | 20 L | 4.1 I + 10.25 T* | 5 $\mu\text{L}$ + 27(10 $\mu\text{L}$) | Ref. 27 | Full 1 |
| IC/ Tctex1 v. LC8 | 107.5 L | 20 I + 50 T* | 3 $\mu\text{L}$ + 24(10 $\mu\text{L}$) | Ref. 27 | Full 2 |
| IC/ Tctex1 v. LC8 | 130 L | 31 I + 77.5 T* | 3 $\mu\text{L}$ + 27(10 $\mu\text{L}$) | Ref. 27 | Full 2 |
| IC/ Tctex1 v. LC8 | 130 L | 31 I + 77.5 T* | 3 $\mu\text{L}$ + 27(10 $\mu\text{L}$) | Ref. 27 | Full 2 |
| IC/ Tctex1 v. LC8 | 223.5 L | 15 I + 37.5 T* | 3 $\mu\text{L}$ + 19(7 $\mu\text{L}$) | Ref. 27 | Full 1 |
| IC/ Tctex1 v. LC8 | 205 L | 23 I + 57.5 T* | 2.5 $\mu\text{L}$ + 29(5 $\mu\text{L}$) | Ref. 27 | Full 1 |
| Tctex1 v. IC | 500 I | 50 T | 2 $\mu\text{L}$ + 23(10 $\mu\text{L}$) | New | T prior |
| Tctex1 v. IC | 500 I | 25 T | 2 $\mu\text{L}$ + 27(10 $\mu\text{L}$) | New | T prior |
| Tctex1 v. IC | 500 I | 12.5 T | 2 $\mu\text{L}$ + 55(5 $\mu\text{L}$) | New | T prior |
| LC8 v. IC | 500 I | 50 L | 2 $\mu\text{L}$ + 27(10 $\mu\text{L}$) | New | L prior |
| LC8 v. IC | 500 I | 25 L | 2 $\mu\text{L}$ + 23(10 $\mu\text{L}$) | New | L prior |
| LC8 v. IC | 500 I | 12.5 L | 2 $\mu\text{L}$ + 55(5 $\mu\text{L}$) | New | L prior |
| LC8/Tctex1 v. IC | 800 I | 30 L + 30 T | 2 $\mu\text{L}$ + 27(5 $\mu\text{L}$) | New | Full 2 |
| LC8/Tctex1 v. IC | 400 I | 30 L + 7.5 T | 2 $\mu\text{L}$ + 20(5 $\mu\text{L}$) + 18(10 $\mu\text{L}$) | New | Full 1 |
| LC8/Tctex1 v. IC | 400 I | 7.5 L + 30 T | 2 $\mu\text{L}$ + 20(5 $\mu\text{L}$) + 18(10 $\mu\text{L}$) | New | Full 2 |
| LC8/Tctex1 v. IC | 160 I | 12 L + 12 T | 2 $\mu\text{L}$ + 27(10 $\mu\text{L}$) | New | Full 1 |
| Dissociation | 50 I + 25 L + 25 T | Buffer | 2 $\mu\text{L}$ + 8(30 $\mu\text{L}$) | New | Full 1 |
| Dissociation | 50 I + 25 L + 25 T | Buffer | (2 + 10 + 10 + 20 + 20 + 40 + 90 + 90) $\mu\text{L}$ | New | Full 2 |

### Bayesian Inference

To analyze the ITC data, we employed the Bayesian hierarchical ITC method(24), which uses the pocoMC(44) Python package to sample from the posterior distribution. The approach enables joint analysis of multiple isotherms, while treating concentration uncertainties as parameters to be fitted. Furthermore, posteriors from previous runs can be used to inform the priors of consecutive runs, enabling the consecutive analysis of multiple sets of isotherms. For analysis, the isotherms were grouped into the seven involving only Tctex1 and IC, the ten involving only LC8 and IC, and the 22 involving all three species. The two two-component groups were each run separately with 16384 particles and four times as many independent samples. Thermodynamic priors were uniform but constrained such that −5 < ΔΔG < 5 and −15 < ΔΔH < 15 for each direct transition as well as restricting the absolute thermodynamic parameters to −20*(b_j_-1) < ΔG_j_ < 0 and −20*(b_j_-1) < ΔH_j_ < 0, with b_j_ the number of binding partners required to form the respective binding state. Concentrations were modelled using the hierarchical approach(24), employing individual lognormal prior distributions centered around the measured value (see Table 3 for measured concentrations) with an additional parent prior on the relative error in the concentration measurements of each species across all experiments with a uniform prior from 0 to 200%.

The thermodynamic value posteriors of the two-component groups were used as a prior distribution for the three-component runs by fitting a kernel density estimate to the full thermodynamic posterior and drawing initial samples for the consecutive runs directly from the posterior samples. The remaining 22 isotherms were divided into two evenly split sets of eleven isotherms for computational efficiency. The Bayesian inference process was applied to each of these groups individually using three replicate runs of 4096 particles and eight times as many independent samples. The combined posteriors of these replicate runs were then used to inform the prior of a consecutive Bayesian run of the other eleven isotherms using identical parameters, resulting in posterior samples from two separate analysis branches.

To further refine the final sampling, we found and combined pairs of samples between the posteriors of these two branches that are close in thermodynamic values, combining the respective sample pairs to form a complete sample for the full set of 22 isotherms, as described in Ref. (45). The combined samples with the best fit of all 22 isotherms as measured by posterior probability, were used as starting points for a final joint run of all three-component isotherms using two replicates of 1024 particles and four times as many independent samples. This final run used the same prior distribution as the two analysis branches before, i.e., the thermodynamic value posteriors of the two-component groups were used as a prior distribution.

### Reported Significant Figures

We determined the number of significant figures we report in our tables from the Bayesian fits based on the half-widths of the 95% confidence intervals. Specifically, the number of decimal places reported is equal to one order of magnitude smaller than the order of magnitude of the 95% confidence interval half-width. For example, the 95% CI for T_I_ ΔH is (−7.07, −6.45) with a half width of 0.31, which is the 10^−1^ order of magnitude, so the significant digits are shown to the 10^−2^ order of magnitude, i.e. −6.76 kcal/mol.

## Supporting information

Supplemental Tables 1-4 and Figures 1-25. Short supplemental discussion for Figure S2.

## Acknowledgments

The authors are grateful for support from the National Institutes of Health under grant GM141733 to E.J.B. and D.M.Z. and from the National Science Foundation under grant MCB 2119837 to D.M.Z. The research reported in this publication used computational infrastructure supported by the Office of Research Infrastructure Programs, Office of the Director, of the National Institutes of Health under award number S10OD034224. The content is solely the responsibility of the authors and does not necessarily represent the official views of the National Institutes of Health.

