## Supplemental Tables 1-4 and Figures 1-25. Short supplemental discussion for Figure S2. for "Walking the Tightrope: Balancing Opposing Cooperativities as an Operating Principle in Dynein Assembly"

Dueling Cooperativity in Dynein Assembly Unveiled via Hierarchical Bayesian Integration of Dozens of Isotherms and a Seven-State Model

Douglas R. Walker^1†^, Lisa Otten^2†^, Mukhtar O. Idris^1^, Brittany Lasher^1^, Daniel M. Zuckerman*^2^, and Elisar J. Barbar*^1^

1. Dept. of Biochemistry and Biophysics, Oregon State University, Corvallis, OR, US.

2. Dept. of Biomedical Engineering, School of Medicine, Oregon Health and Science University, Portland, OR, US.

^†^These authors contributed equally.

**Supplemental Information:**

Table S1. Measured thermodynamic values (in kcal/mol) and 95% confidence intervals from Bayesian inference of two-protein isotherms. As an example for free-energy notation, ΔG(T_II_) = (G(T_II_) - G(T_I_)) and ΔΔG(T_II_) = ΔG(T_II_) - G(T_I_).

| **Species** | **ΔG** | **95% CI** | **ΔΔG** | **95% CI** | **ΔH** | **95% CI** | **ΔΔH** | **95% CI** |
| --- | --- | --- | --- | --- | --- | --- | --- | --- |
| T_I_ | -6.38 | (-6.60, -6.18) | - | - | -5.3 | (-7.2, -3.8) | - | - |
| T_II_ | -6.58 | (-6.76, -6.39) | -0.19 | (-0.33, -0.03) | -4.1 | (-5.6, -2.9) | 1.2 | (0.2, 2.4) |
| L_I_ | -6.710 | (-6.801, -6.611) | - | - | -12.7 | (-14.0, -11.0) | - | - |
| L_II_ | -6.972 | (-7.036, -6.887) | -0.263 | (-0.347, -0.171) | -13.2 | (-14.5, -11.6) | -0.51 | (-1.08, -0.02) |

Table S2. Absolute thermodynamic values (in kcal/mol) and 95% confidence interval ranges for each state as depicted in Figure 5b.

| Species | G | 95% CI | H | 95 % CI |
| --- | --- | --- | --- | --- |
| T_I_ | -6.645 | (-6.695, -6.594) | -6.76 | (-7.07, -6.45) |
| L_I_ | -6.804 | (-6.845, -6.758) | -14.75 | (-15.23, -14.29) |
| T_II_ | -13.385 | (-13.439, -13.329) | -11.73 | (-12.24, -11.22) |
| T_II_ (null) | -13.29 | (-13.39, -13.19) | -13.53 | (-14.13, -12.90) |
| L_II_ | -13.870 | (-13.917, -13.819) | -29.81 | (-30.78, -28.89) |
| L_II_ (null) | -13.608 | (-13.690, -13.516) | -29.51 | (-30.45, -28.58) |
| T_I_L_I_ | -13.16 | (-13.54, -12.54) | -19.4 | (-21.0, -17.9) |
| T_I_L_I_ (null) | -13.449 | (-13.516, -13.367) | -21.52 | (-22.21, -20.81) |
| T_II_L_II_ | -24.997 | (-25.094, -24.895) | -37.5 | (-38.6, -36.4) |
| T_II_L_II_ (null, ΔΔ_1_) | -26.90 | (-27.03, -26.73) | -43.0 | (-44.4, -41.6) |
| T_II_L_II_ (null, ΔΔ_2_) | -27.255 | (-27.342, -27.160) | -41.5 | (-42.9, -40.2) |
| T_II_L_II_ (null, ΔΔ_3_) | -26.97 | (-27.35, -26.35) | -39.5 | (-41.2, 37.6) |
| T_II_L_II_ (null, ΔΔ_4_) | -26.3 | (-27.1, -25.1) | -38.9 | (-41.9, -35.8) |
| T_II_L_II_ (null, ΔΔ_5_) | -26.68 | (-27.34, -25.44) | -37.4 | (-40.2, -34.4) |

Table S3. Measured entropies (TΔS in kcal/mol) and 95% confidence interval ranges for all transitions and cooperativities based on all isotherms.

| **Transition** | **TΔS** | **95% CI** | **Δ(TΔS)** | **95% CI** |
| --- | --- | --- | --- | --- |
| T_I_ | -0.12 | (-0.45, 0.22) | - | - |
| T_II_ | 1.77 | (1.44, 2.10) | 1.89 | (1.44, 2.31) |
| L_I_ | -7.95 | (-8.42, -7.50) | - | - |
| L_II_ | -7.99 | (-8.52, -7.50) | -0.04 | (-0.40, 0.31) |
| T_I_ -> T_I_L_I_ | -6.2 | (-7.7, -4.8) | 1.8 | (0.2, 3.1) |
| L_I_ -> T_I_L_I_ | 1.7 | (0.0, 3.0) |  |  |
| T_II_ -> T_II_L_II_ | -14.18 | (-15.00, -13.40) | ΔΔ_3_ =  -0.0 | (-1.3, 1.5) |
| T_I_L_I_ -> T_II_L_II_ | -6.2 | (-7.7, -4.6) |  |  |
| L_II_ -> T_II_L_II_ | 3.41 | (2.92, 3.92) |  |  |

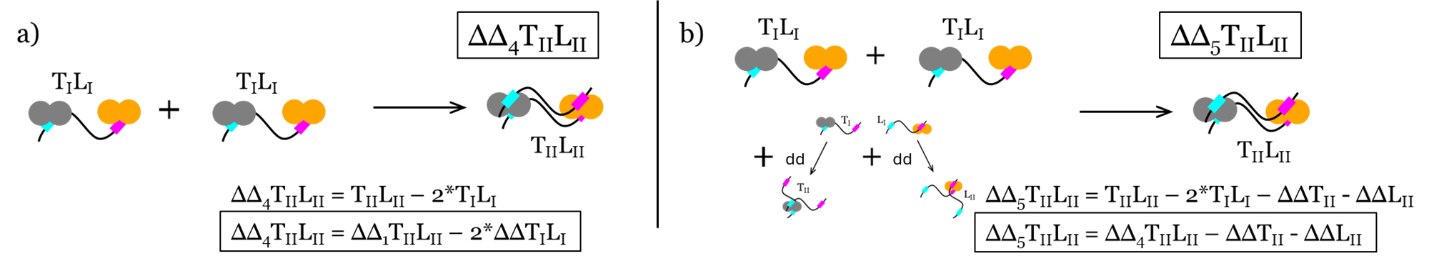
Table S4. Measured thermodynamic cooperativities (in kcal/mol) and 95% confidence interval ranges for the different perspectives on cooperativity in the fully bound state based on all isotherms.

| **Scheme** | **ΔΔG** | **95% CI** | **ΔΔH** | **95% CI** | **Δ(TΔS)** | **95% CI** |
| --- | --- | --- | --- | --- | --- | --- |
| ΔΔ_1_ | 1.901 | (1.798, 1.993) | 5.51 | (4.92, 6.16) | 3.61 | (2.96, 4.33) |
| ΔΔ_2_ | 2.258 | (2.178, 2.332) | 4.02 | (3.42, 4.66) | 1.76 | (1.16, 2.41) |
| ΔΔ_3_ | 1.97 | (1.40, 2.31) | 1.9 | (0.4, 3.3) | -0.0 | (-1.3, 1.5) |
| ΔΔ_4_ | 1.33 | (0.11, 2.03) | 1.4 | (-1.6, 4.2) | 0.0 | (-2.6, 3.2) |
| ΔΔ_5_ | 1.68 | (0.49, 2.39) | -0.1 | (-3.0, 2.5) | -1.8 | (-4.3, 1.2) |

Figure S1. Two additional perspectives from which to consider the “constituent parts” of the trivalent T_II_L_II_. The distributions of these cooperativities are tabulated in Table S4 but were not shown in the main manuscript Fig. 5 due to the broadness of these distributions. Readers may note that the source of this broadness is the two-fold inclusion of the T_I_L_I_ state in their calculation.

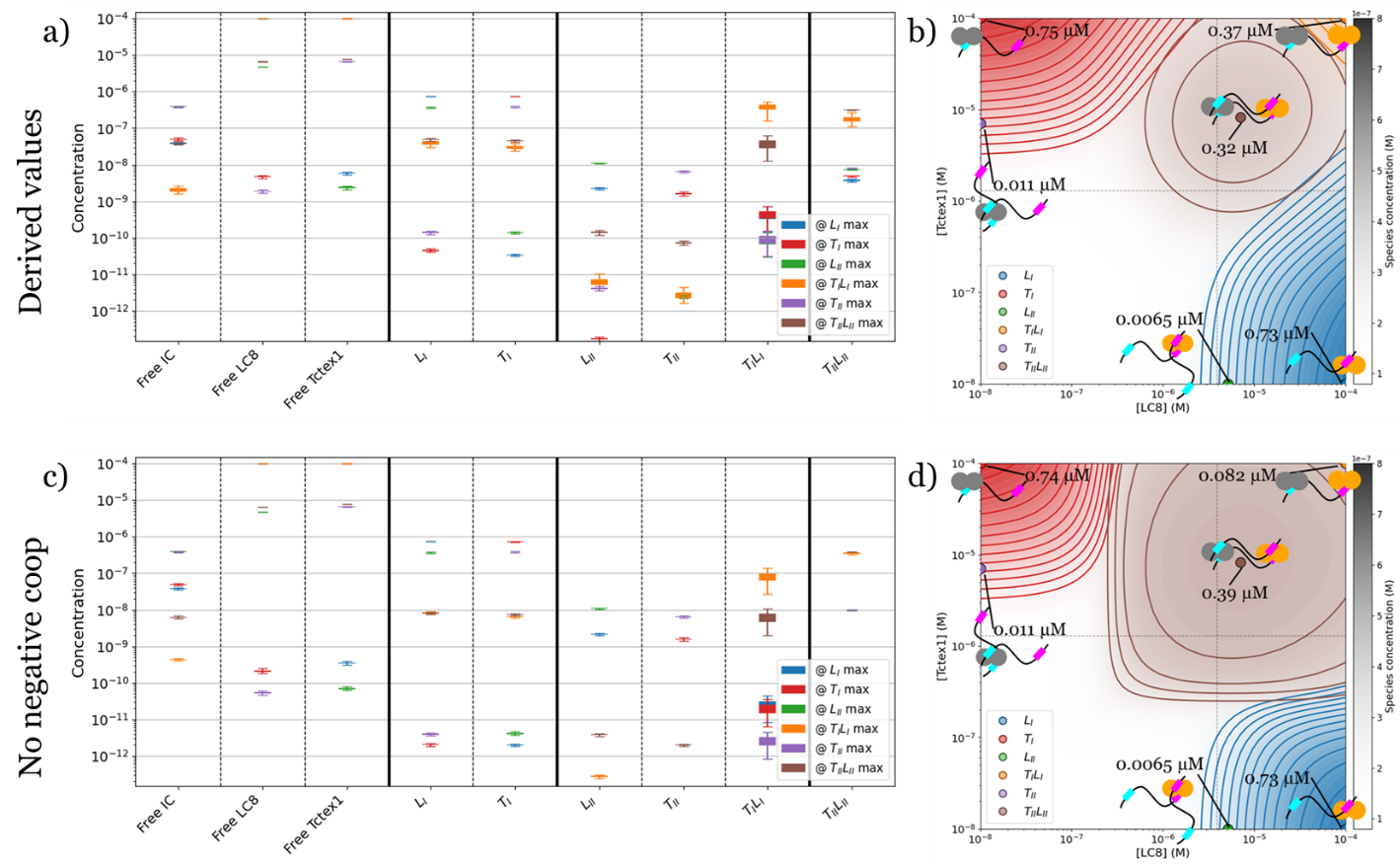
 Figure S2. Effects of cooperativity on state populations. State-specific concentrations at different total concentrations of LC8 and Tctex1 are compared to a system with no cooperativity. a) Concentrations and associated errors at seven different concentrations of LC8 and Tctex1 based on Bayesian-inferred thermodynamics. The seven points were chosen as the combinations which maximize one of the seven states over the matrix of LC8 and Tctex1 concentrations examined in panel b. b) Concentration contours for each of the seven states as [LC8] and [Tctex1] vary between 10 nM and 0.1 mM. c & d) Same as a & b for the case in which the energy of fully bound state is adjusted to remove cooperativity, specifically it is set as the sum of T_II_ and L_II_: G_null_(T_II_L_II_) = G(T_II_) + G(L_II_). In all panels, [IC] = 0.8 µM. Dotted lines in b and d denote where [LC8] = [IC] or where [Tctex1] = [IC].

Although less explicit than at the simulations shown in Fig. 6, Fig. S2 also illustrates the accessibility of T_I_L_I_ at elevated concentrations of LC8 and Tctex1 and that this state would not be accessible without the negative cooperativity of the system. This simulation represents what we believe is the lower limit on that accessibility because while this simulation utilized previously reported intra-cellular IC concentrations, we cannot account for increased local concentrations throughout a heterogeneous cellular environment, nor for the effects of crowding on equilibrium, both of which should result in pushes in equilibrium toward the upper-right of the state concentration plot in Fig. S2b.

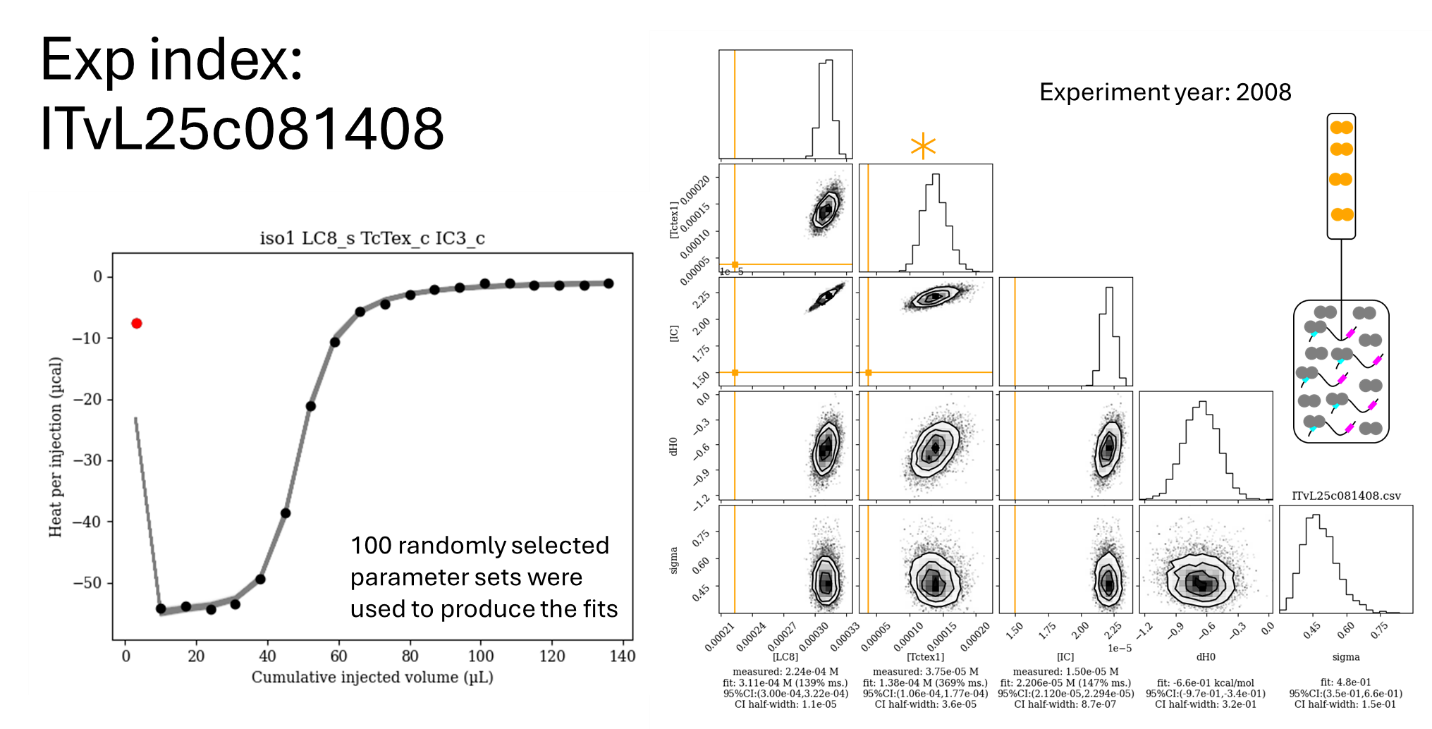
Figure S3. Isotherm ITvL25c081408 with the distributions of the fits of the concentrations and other nuisance parameters associated with the given isotherm. 100 random parameter sets from the posterior distribution were selected for drawing the fits on the isotherm. As shown in the cartoon, LC8 was titrated into a mixture of IC and Tctex1 for this experiment. The red points in the isotherm indicate injections that were disregarded in the fit. Orange bars on the corner plot correspond to values input as “measured”. The asterisk denotes that the Tctex1 concentration was not recorded for this experiment but was referred to as being ~5 times that of the IC concentration.

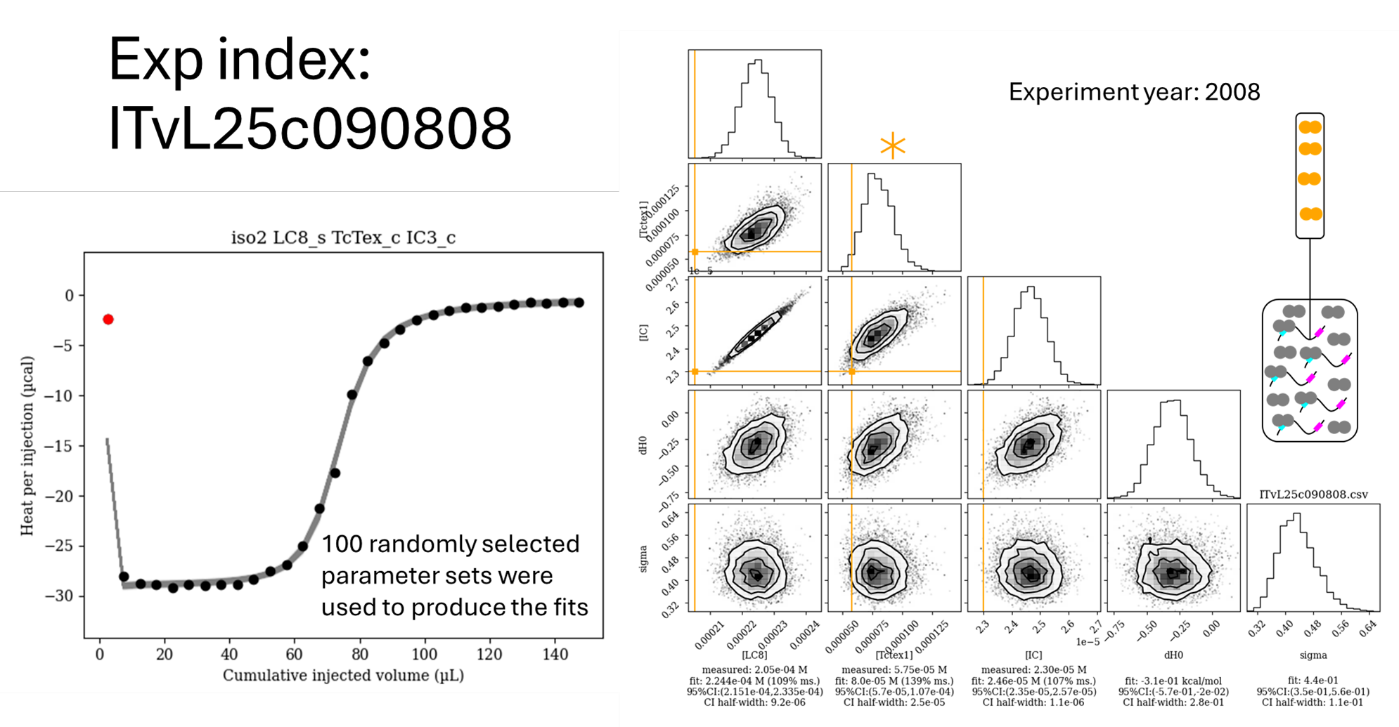

Figure S4. Isotherm ITvL25c090808 with the distributions of the fits of the concentrations and other nuisance parameters associated with the given isotherm. 100 random parameter sets from the posterior distribution were selected for drawing the fits on the isotherm. As shown in the cartoon, LC8 was titrated into a mixture of IC and Tctex1 for this experiment. The red points in the isotherm indicate injections that were disregarded in the fit. Orange bars on the corner plot correspond to values input as “measured”. The asterisk denotes that the Tctex1 concentration was not recorded for this experiment but was referred to as being ~5 times that of the IC concentration.

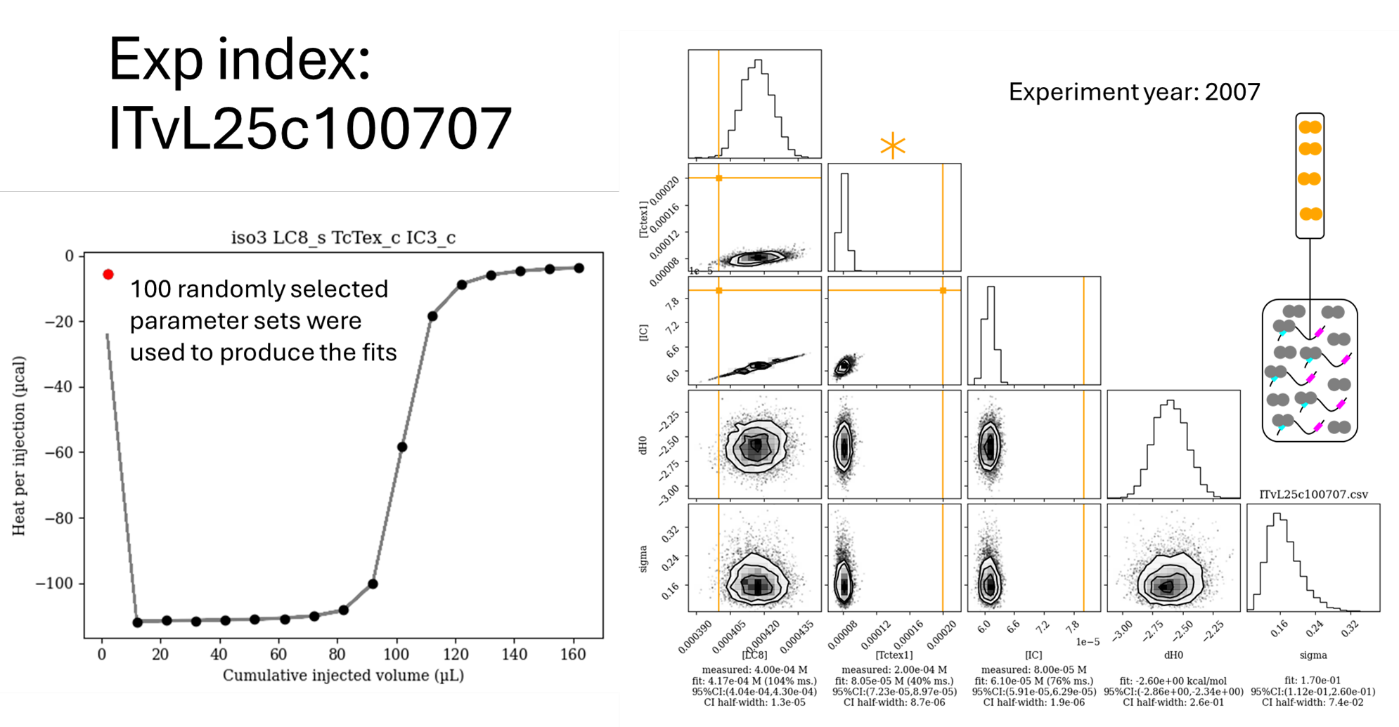

Figure S5. Isotherm ITvL25c100707 with the distributions of the fits of the concentrations and other nuisance parameters associated with the given isotherm. 100 random parameter sets from the posterior distribution were selected for drawing the fits on the isotherm. As shown in the cartoon, LC8 was titrated into a mixture of IC and Tctex1 for this experiment. The red points in the isotherm indicate injections that were disregarded in the fit. Orange bars on the corner plot correspond to values input as “measured”. The asterisk denotes that the Tctex1 concentration was not recorded for this experiment but was referred to as being ~5 times that of the IC concentration.

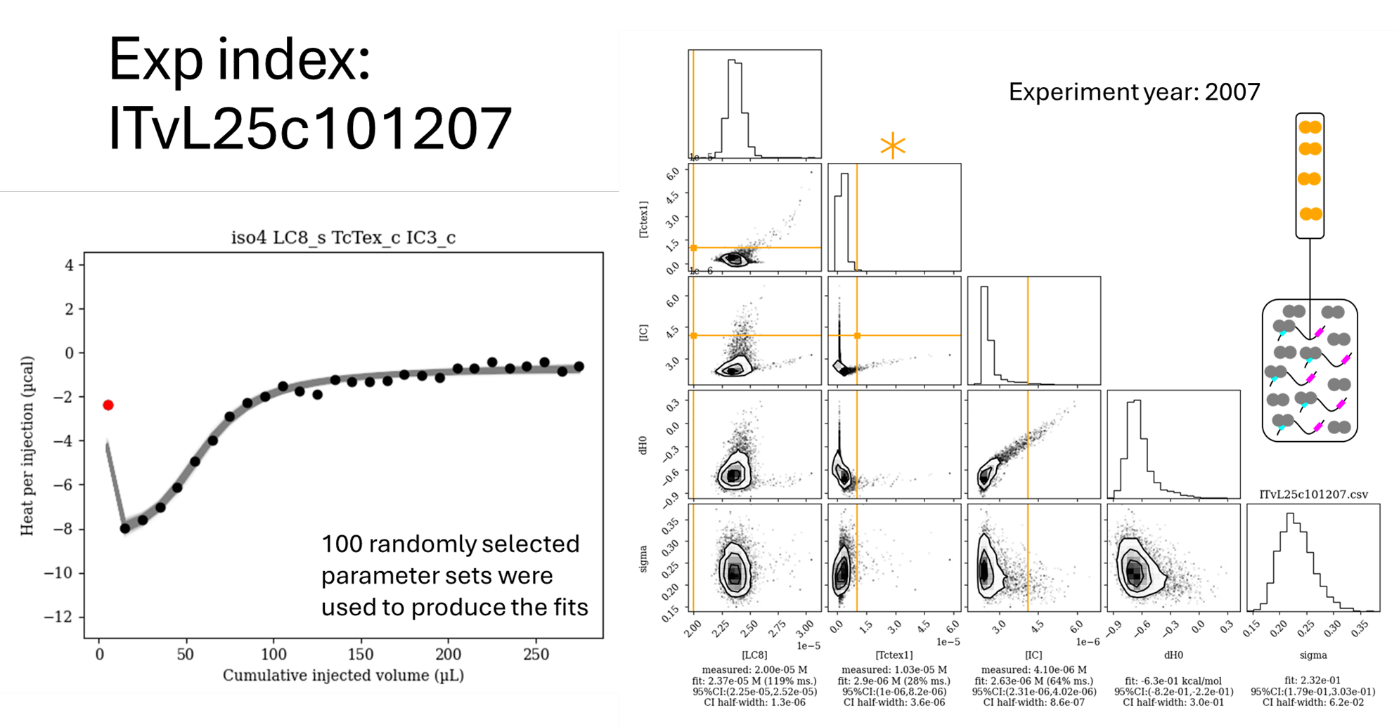

Figure S6. Isotherm ITvL25c101207 with the distributions of the fits of the concentrations and other nuisance parameters associated with the given isotherm. 100 random parameter sets from the posterior distribution were selected for drawing the fits on the isotherm. As shown in the cartoon, LC8 was titrated into a mixture of IC and Tctex1 for this experiment. The red points in the isotherm indicate injections that were disregarded in the fit. Orange bars on the corner plot correspond to values input as “measured”. The asterisk denotes that the Tctex1 concentration was not recorded for this experiment but was referred to as being ~5 times that of the IC concentration.

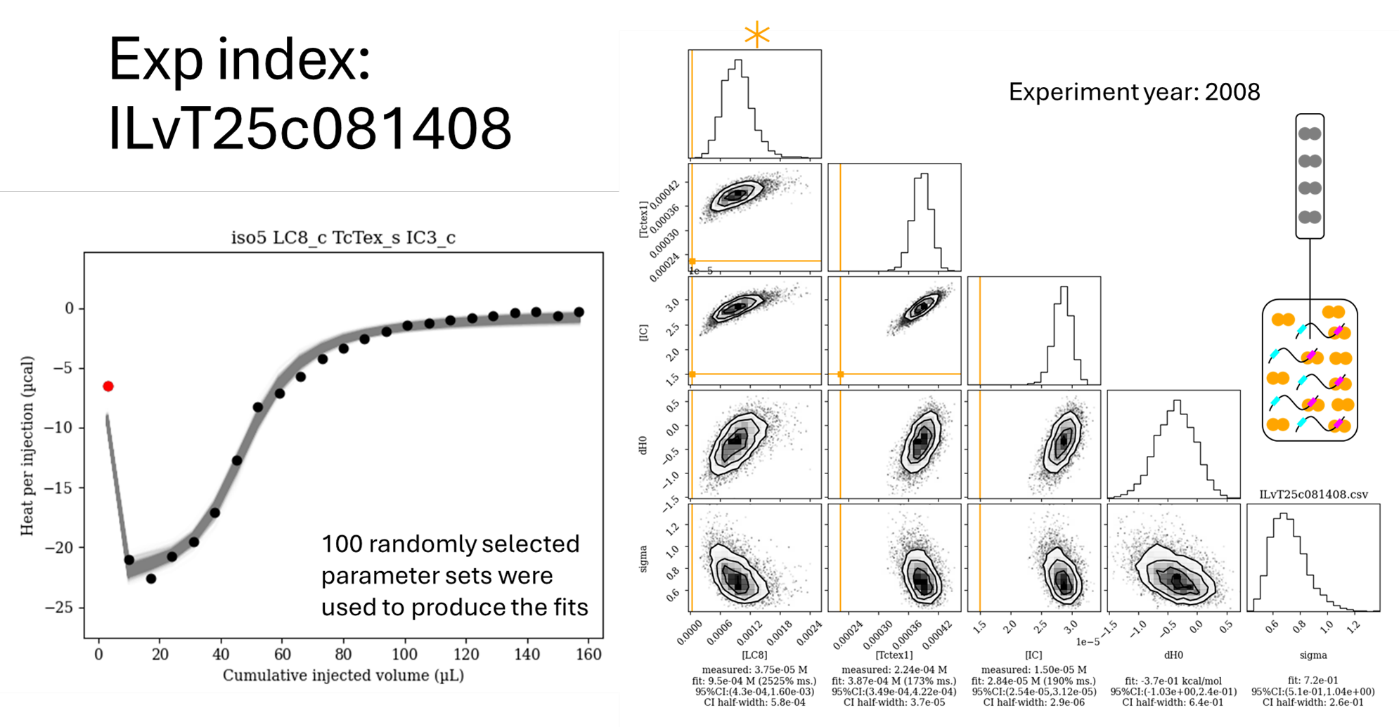

Figure S7. Isotherm ILvT25c081408 with the distributions of the fits of the concentrations and other nuisance parameters associated with the given isotherm. 100 random parameter sets from the posterior distribution were selected for drawing the fits on the isotherm. As shown in the cartoon, Tctex1 was titrated into a mixture of IC and LC8 for this experiment. The red points in the isotherm indicate injections that were disregarded in the fit. Orange bars on the corner plot correspond to values input as “measured”. The asterisk denotes that the LC8 concentration was not recorded for this experiment but was referred to as being ~5 times that of the IC concentration.

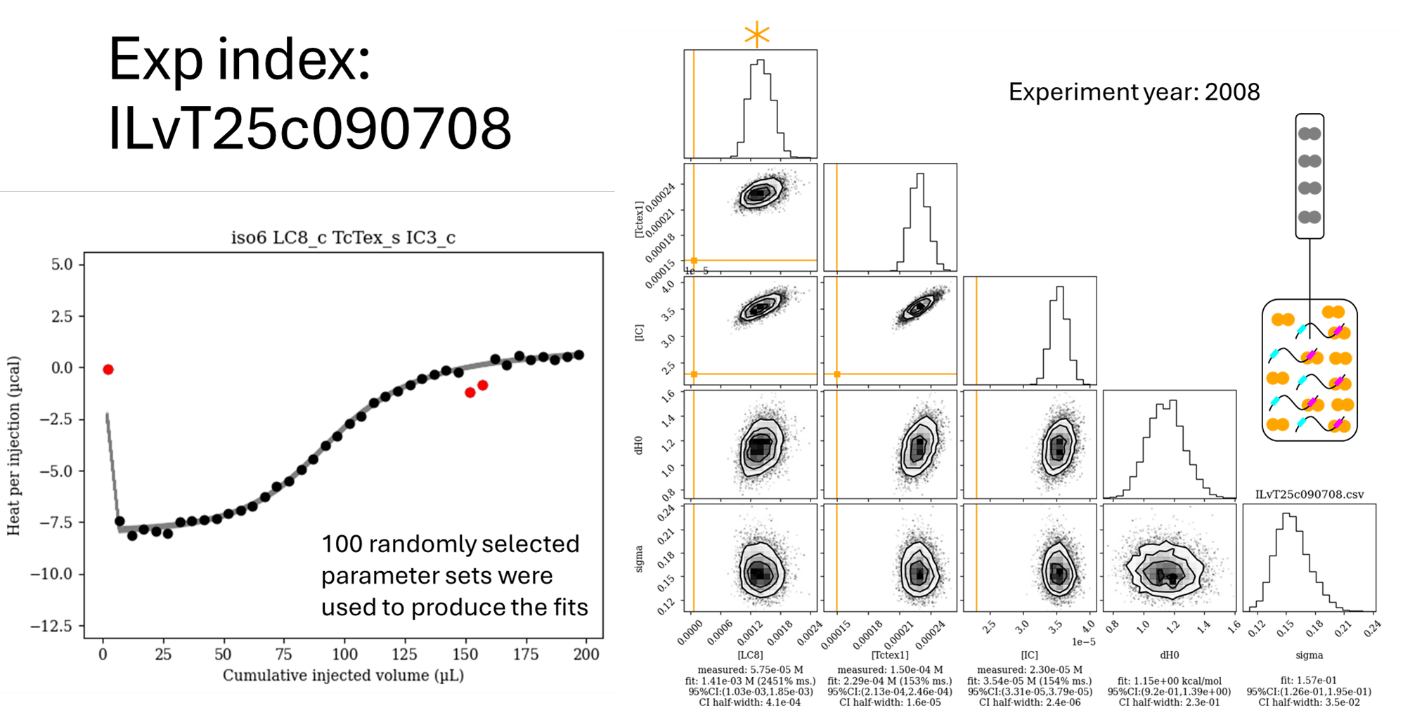

Figure S8. Isotherm ILvT25c090708 with the distributions of the fits of the concentrations and other nuisance parameters associated with the given isotherm. 100 random parameter sets from the posterior distribution were selected for drawing the fits on the isotherm. As shown in the cartoon, Tctex1 was titrated into a mixture of IC and LC8 for this experiment. The red points in the isotherm indicate injections that were disregarded in the fit. Orange bars on the corner plot correspond to values input as “measured”. The asterisk denotes that the LC8 concentration was not recorded for this experiment but was referred to as being ~5 times that of the IC concentration.

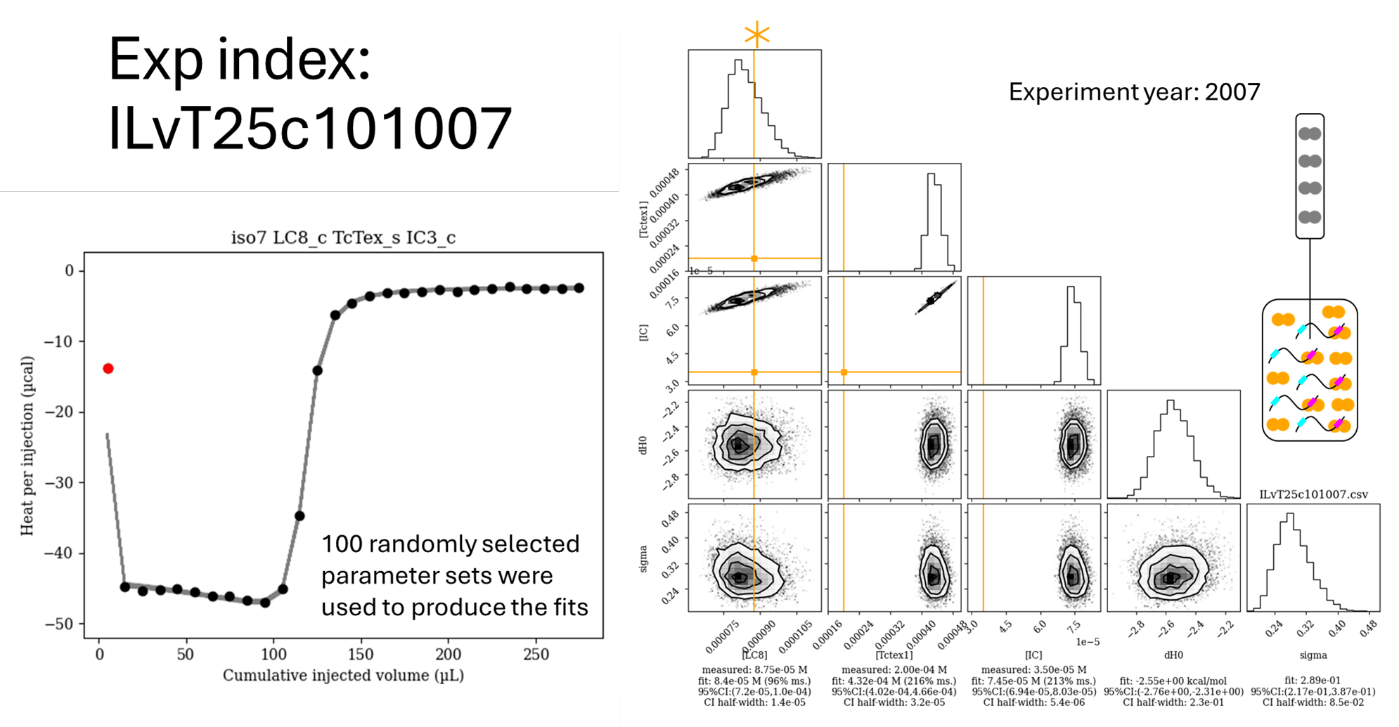

Figure S9. Isotherm ILvT25c101007 with the distributions of the fits of the concentrations and other nuisance parameters associated with the given isotherm. 100 random parameter sets from the posterior distribution were selected for drawing the fits on the isotherm. As shown in the cartoon, Tctex1 was titrated into a mixture of IC and LC8 for this experiment. The red points in the isotherm indicate injections that were disregarded in the fit. Orange bars on the corner plot correspond to values input as “measured”. The asterisk denotes that the LC8 concentration was not recorded for this experiment but was referred to as being ~5 times that of the IC concentration.

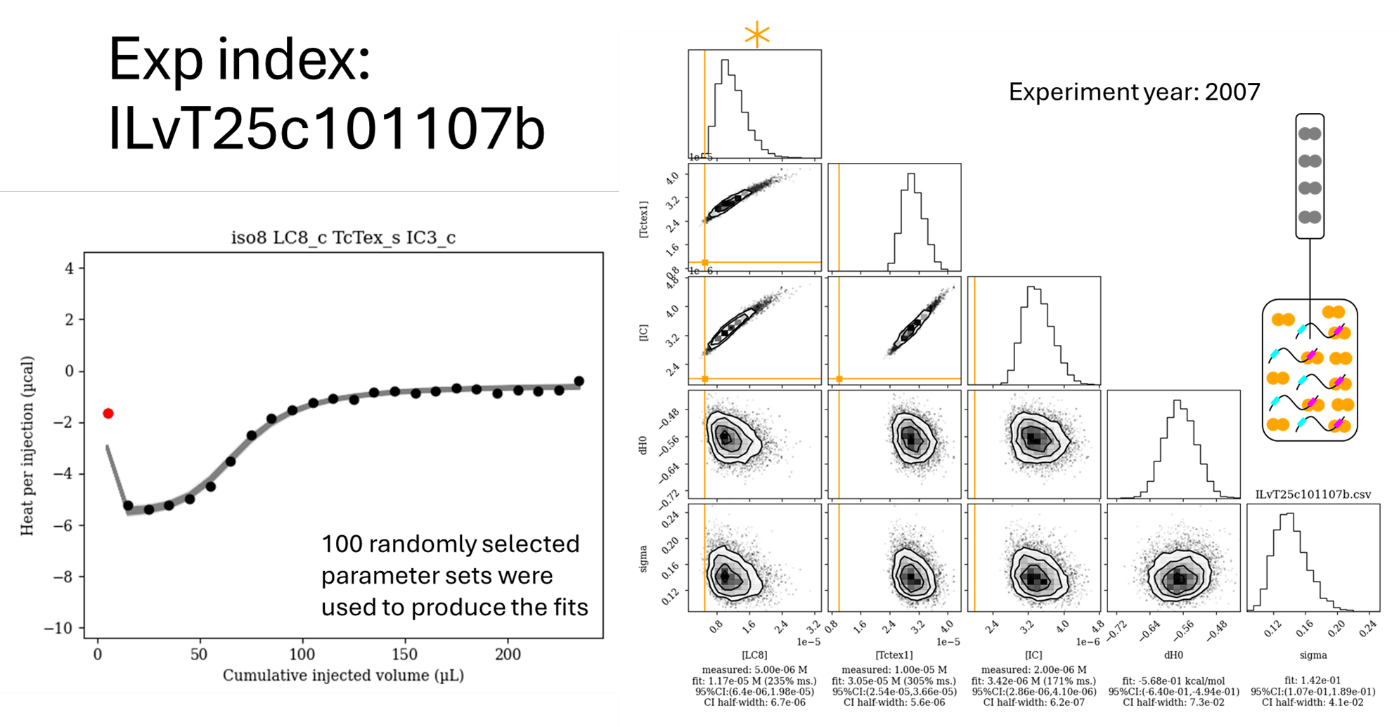

Figure S10. Isotherm ITvL25c101107b with the distributions of the fits of the concentrations and other nuisance parameters associated with the given isotherm. 100 random parameter sets from the posterior distribution were selected for drawing the fits on the isotherm. As shown in the cartoon, Tctex1 was titrated into a mixture of IC and LC8 for this experiment. The red points in the isotherm indicate injections that were disregarded in the fit. Orange bars on the corner plot correspond to values input as “measured”. The asterisk denotes that the LC8 concentration was not recorded for this experiment but was referred to as being ~5 times that of the IC concentration.

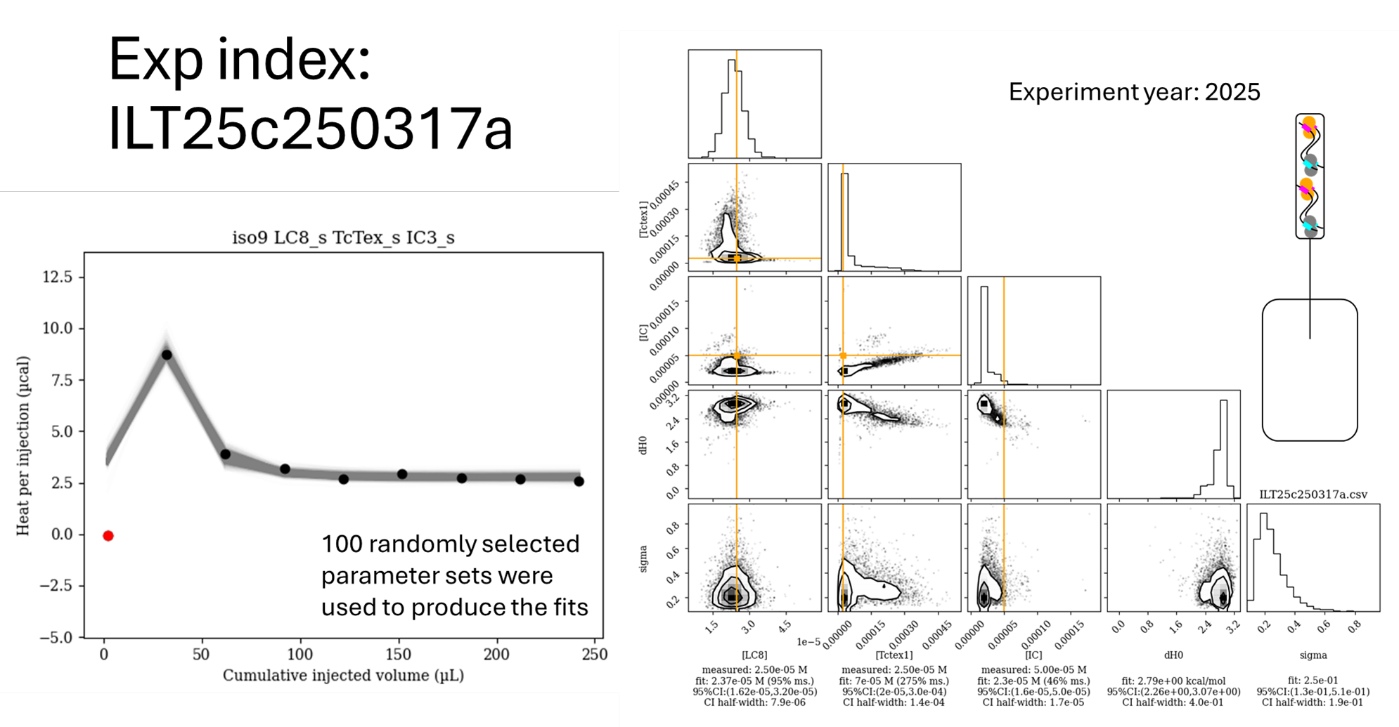

Figure S11. Isotherm ILT25c250317a with the distributions of the fits of the concentrations and other nuisance parameters associated with the given isotherm. 100 random parameter sets from the posterior distribution were selected for drawing the fits on the isotherm. As shown in the cartoon, a mixture of IC, LC8, and Tctex1 was titrated into buffer for this experiment. The red points in the isotherm indicate injections that were disregarded in the fit. Orange bars on the corner plot correspond to values input as “measured”.

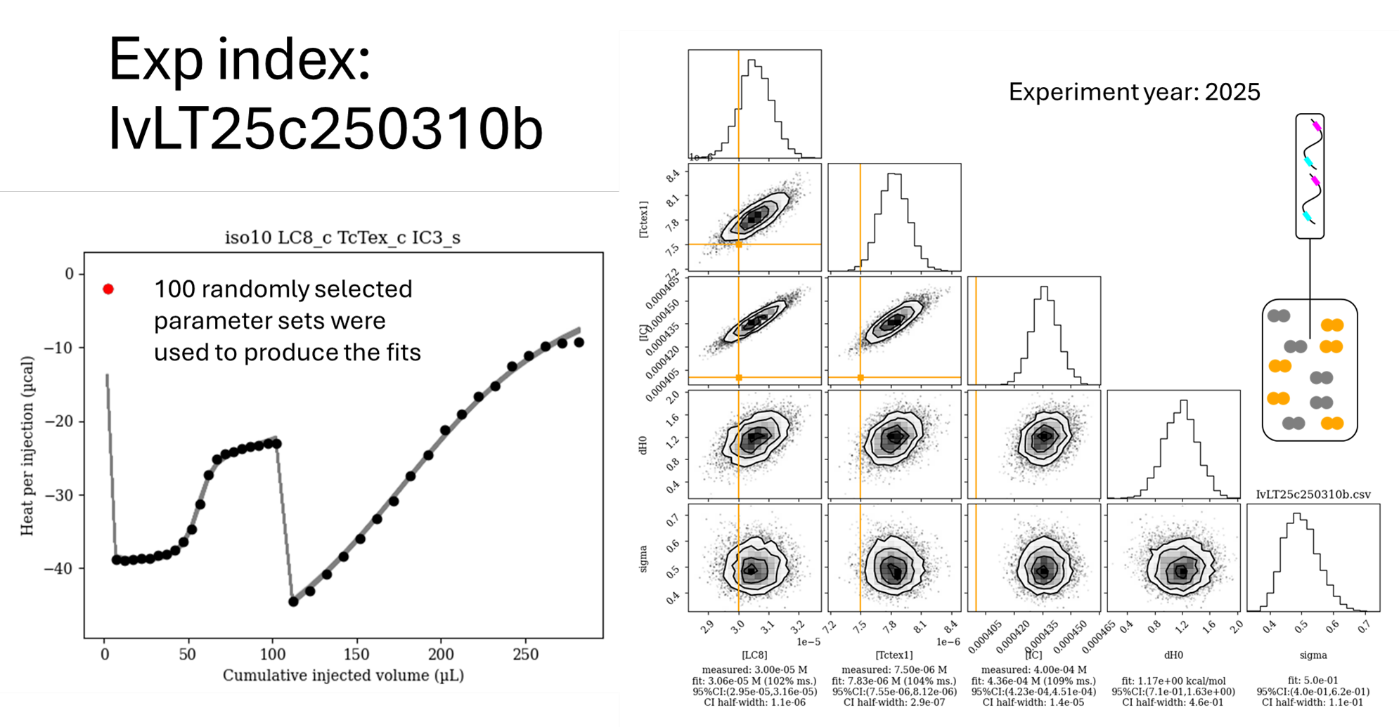

Figure S12. Isotherm IvLT25c250310b with the distributions of the fits of the concentrations and other nuisance parameters associated with the given isotherm. 100 random parameter sets from the posterior distribution were selected for drawing the fits on the isotherm. As shown in the cartoon, IC was titrated into a mixture of LC8 and Tctex1 for this experiment. Notedly, larger injection volumes were used for the second half of the experiment. The red points in the isotherm indicate injections that were disregarded in the fit. Orange bars on the corner plot correspond to values input as “measured”.

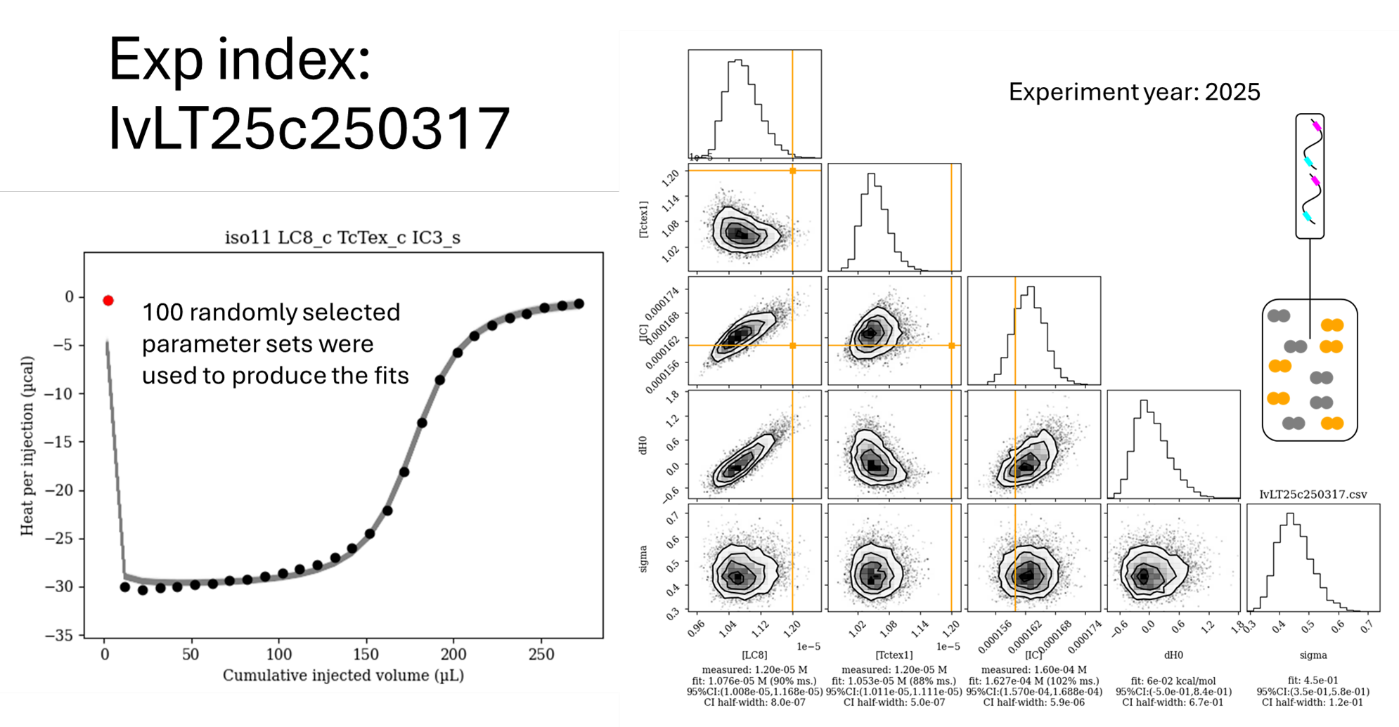

Figure S13. Isotherm IvLT25c250317 with the distributions of the fits of the concentrations and other nuisance parameters associated with the given isotherm. 100 random parameter sets from the posterior distribution were selected for drawing the fits on the isotherm. As shown in the cartoon, IC was titrated into a mixture of LC8 and Tctex1 for this experiment. The red points in the isotherm indicate injections that were disregarded in the fit. Orange bars on the corner plot correspond to values input as “measured”.

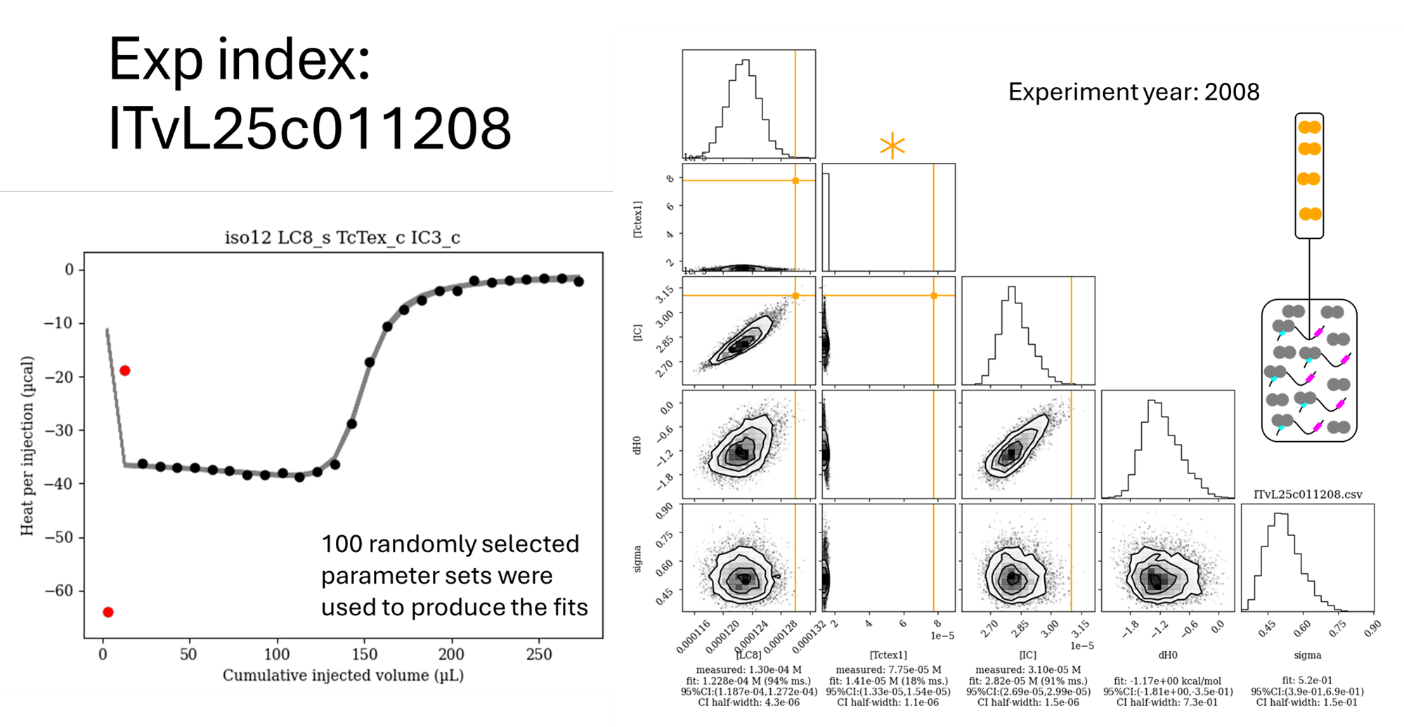

Figure S14. Isotherm ITvL25c011208 with the distributions of the fits of the concentrations and other nuisance parameters associated with the given isotherm. 100 random parameter sets from the posterior distribution were selected for drawing the fits on the isotherm. As shown in the cartoon, LC8 was titrated into a mixture of IC and Tctex1 for this experiment. The red points in the isotherm indicate injections that were disregarded in the fit. Orange bars on the corner plot correspond to values input as “measured”. The asterisk denotes that the Tctex1 concentration was not recorded for this experiment but was referred to as being ~5 times that of the IC concentration.

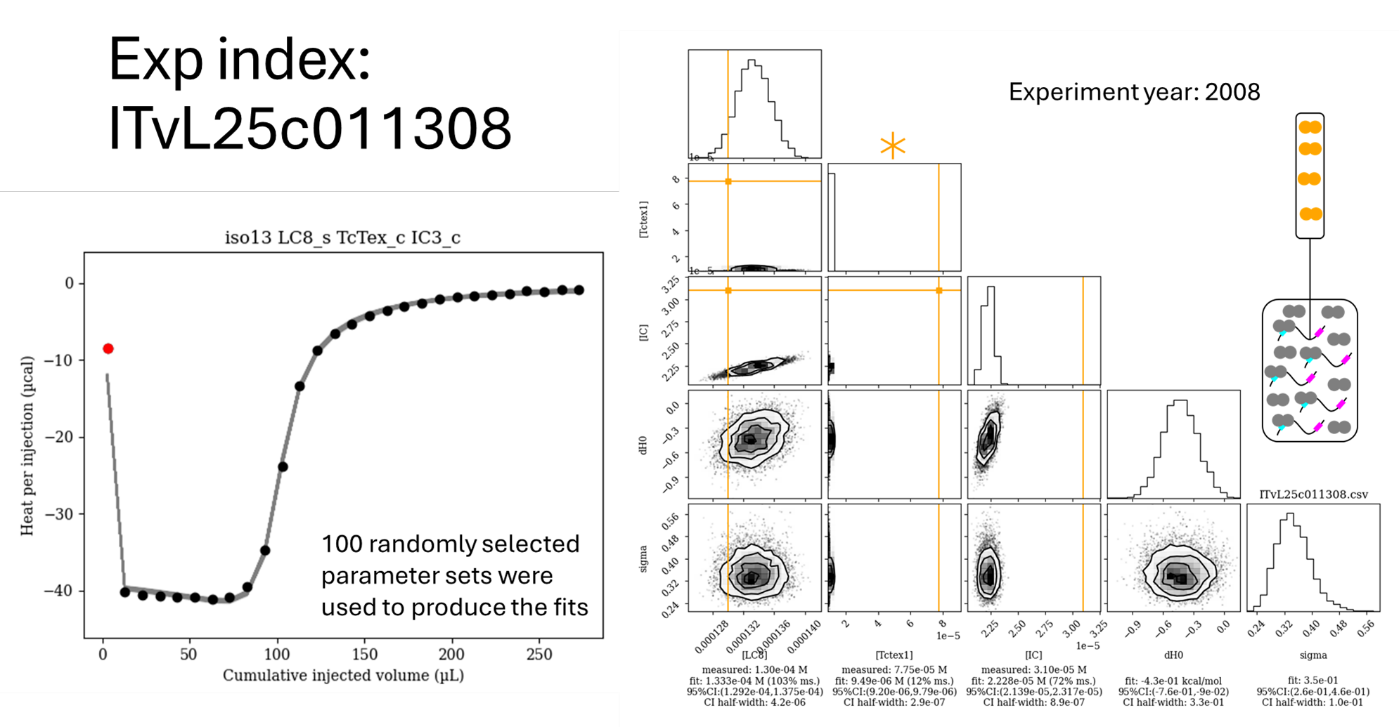

Figure S15. Isotherm ITvL25c011308 with the distributions of the fits of the concentrations and other nuisance parameters associated with the given isotherm. 100 random parameter sets from the posterior distribution were selected for drawing the fits on the isotherm. As shown in the cartoon, LC8 was titrated into a mixture of IC and Tctex1 for this experiment. The red points in the isotherm indicate injections that were disregarded in the fit. Orange bars on the corner plot correspond to values input as “measured”. The asterisk denotes that the Tctex1 concentration was not recorded for this experiment but was referred to as being ~5 times that of the IC concentration.

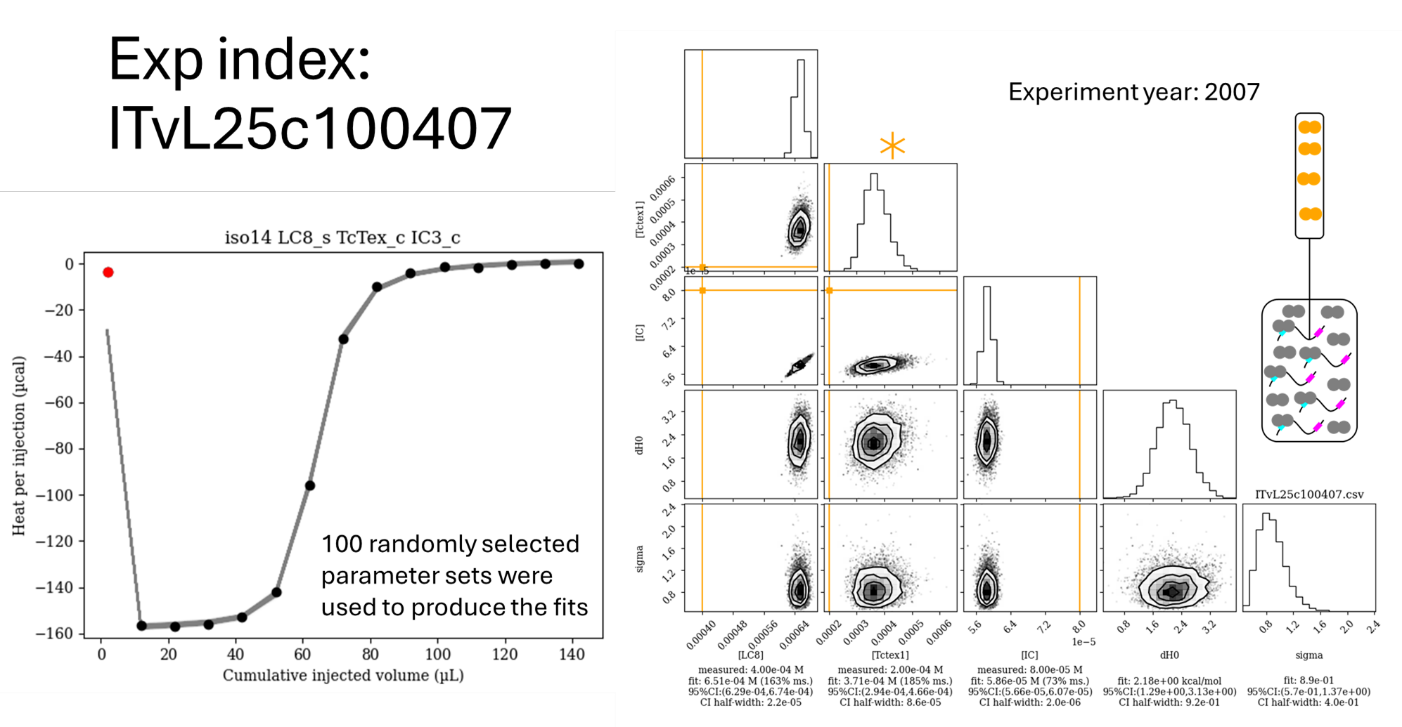

Figure S16. Isotherm ITvL25c100407 with the distributions of the fits of the concentrations and other nuisance parameters associated with the given isotherm. 100 random parameter sets from the posterior distribution were selected for drawing the fits on the isotherm. As shown in the cartoon, LC8 was titrated into a mixture of IC and Tctex1 for this experiment. The red points in the isotherm indicate injections that were disregarded in the fit. Orange bars on the corner plot correspond to values input as “measured”. The asterisk denotes that the Tctex1 concentration was not recorded for this experiment but was referred to as being ~5 times that of the IC concentration.

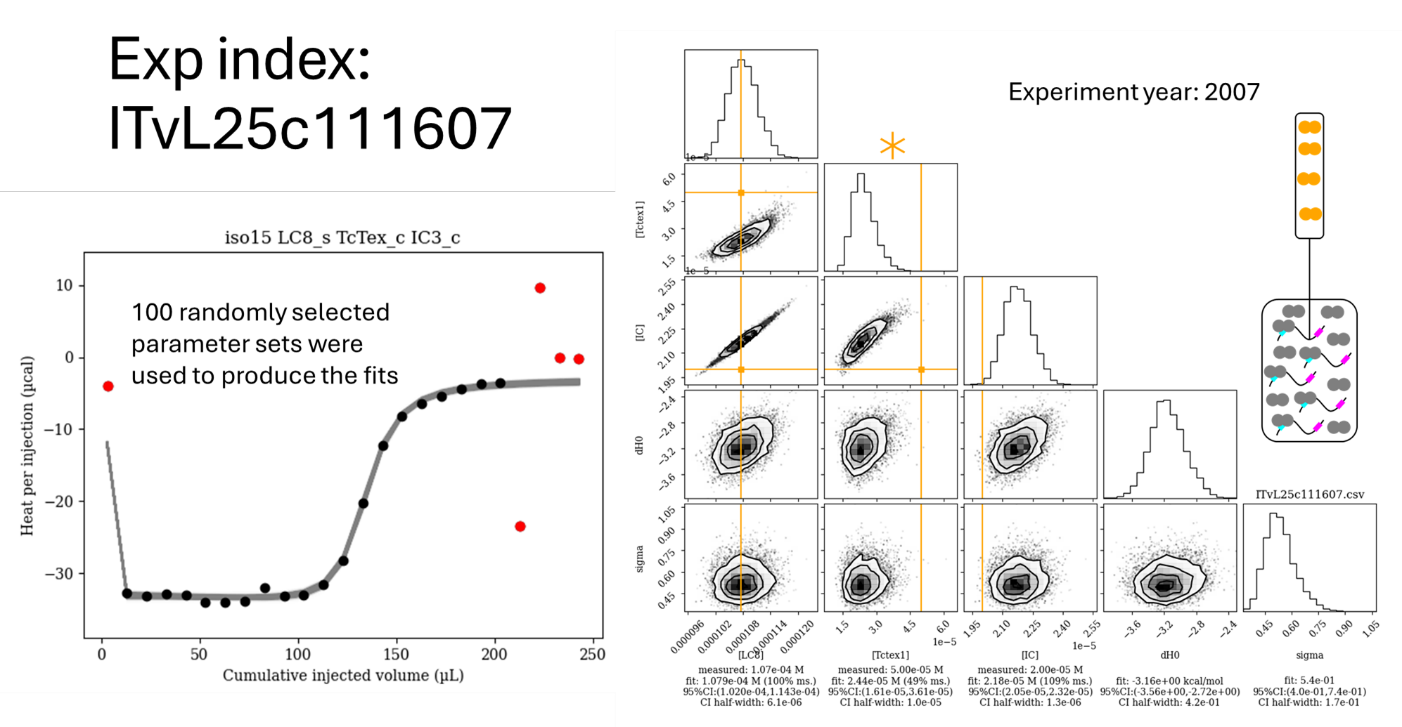

Figure S17. Isotherm ITvL25c111607 with the distributions of the fits of the concentrations and other nuisance parameters associated with the given isotherm. 100 random parameter sets from the posterior distribution were selected for drawing the fits on the isotherm. As shown in the cartoon, LC8 was titrated into a mixture of IC and Tctex1 for this experiment. The red points in the isotherm indicate injections that were disregarded in the fit. Orange bars on the corner plot correspond to values input as “measured”. The asterisk denotes that the Tctex1 concentration was not recorded for this experiment but was referred to as being ~5 times that of the IC concentration.

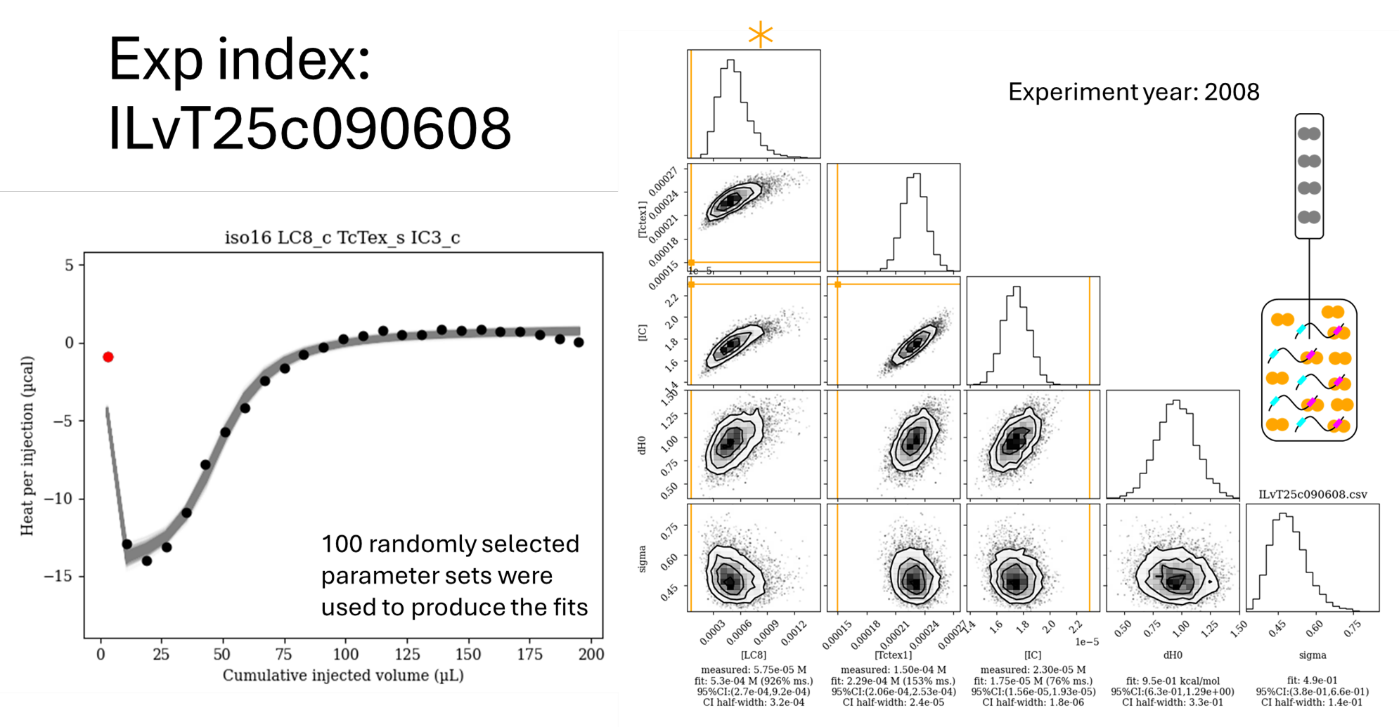

Figure S18. Isotherm ILvT25c090608 with the distributions of the fits of the concentrations and other nuisance parameters associated with the given isotherm. 100 random parameter sets from the posterior distribution were selected for drawing the fits on the isotherm. As shown in the cartoon, Tctex1 was titrated into a mixture of IC and LC8 for this experiment. The red points in the isotherm indicate injections that were disregarded in the fit. Orange bars on the corner plot correspond to values input as “measured”. The asterisk denotes that the LC8 concentration was not recorded for this experiment but was referred to as being ~5 times that of the IC concentration.

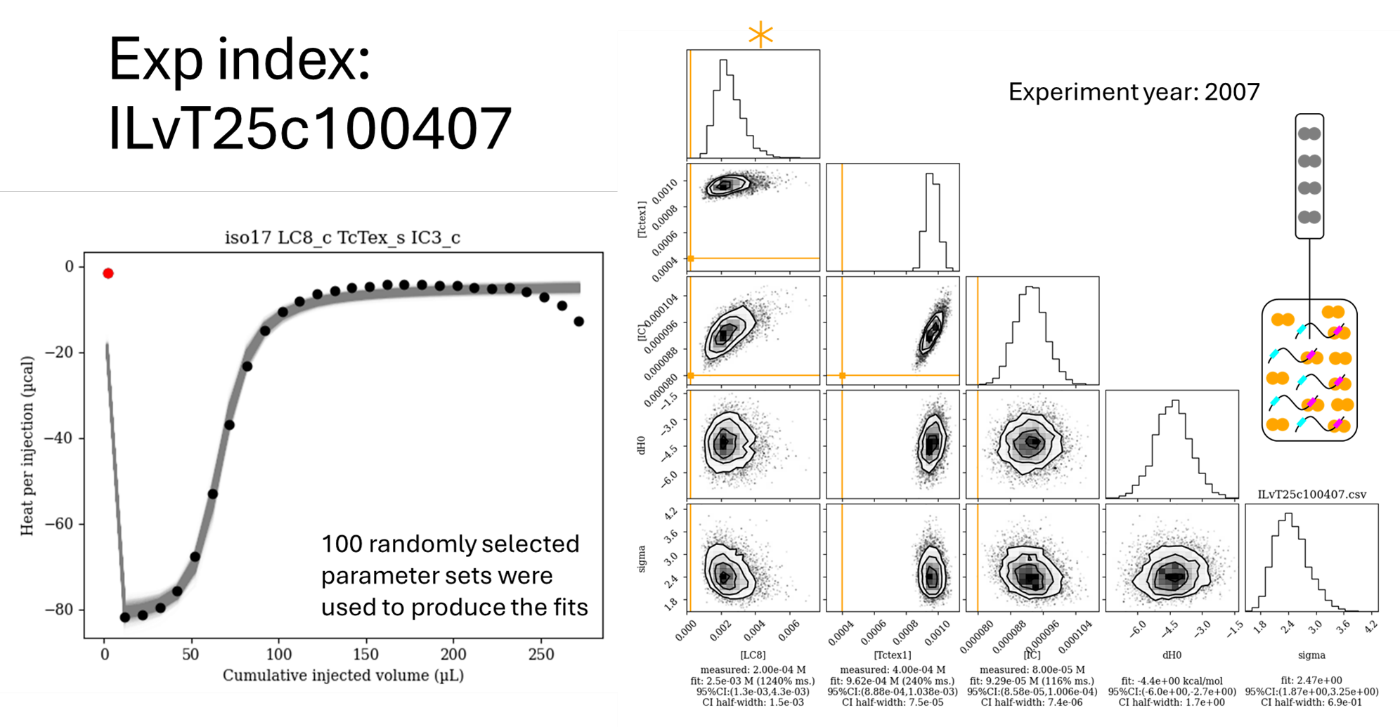

Figure S19. Isotherm ITvL25c100407 with the distributions of the fits of the concentrations and other nuisance parameters associated with the given isotherm. 100 random parameter sets from the posterior distribution were selected for drawing the fits on the isotherm. As shown in the cartoon, Tctex1 was titrated into a mixture of IC and LC8 for this experiment. The red points in the isotherm indicate injections that were disregarded in the fit. Orange bars on the corner plot correspond to values input as “measured”. The asterisk denotes that the LC8 concentration was not recorded for this experiment but was referred to as being ~5 times that of the IC concentration.

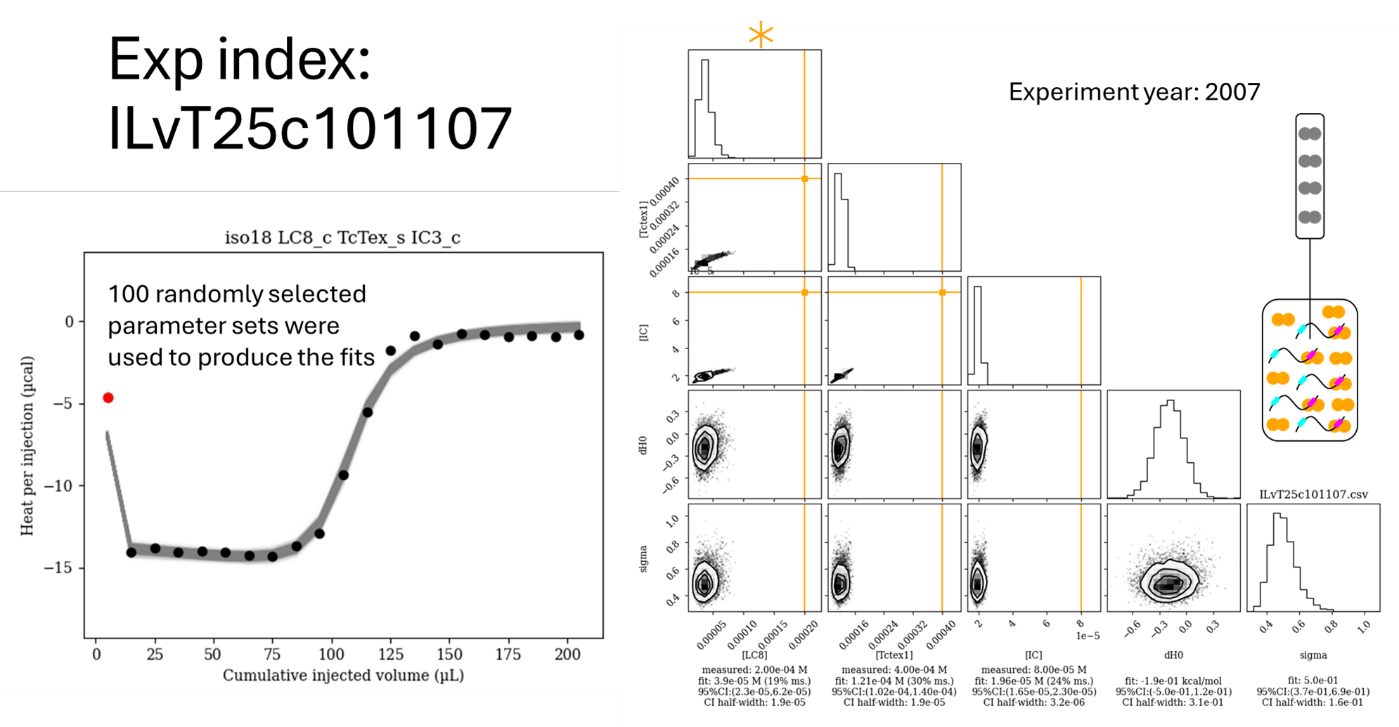

Figure S20. Isotherm ILvT25c101107 with the distributions of the fits of the concentrations and other nuisance parameters associated with the given isotherm. 100 random parameter sets from the posterior distribution were selected for drawing the fits on the isotherm. As shown in the cartoon, Tctex1 was titrated into a mixture of IC and LC8 for this experiment. The red points in the isotherm indicate injections that were disregarded in the fit. Orange bars on the corner plot correspond to values input as “measured”. The asterisk denotes that the LC8 concentration was not recorded for this experiment but was referred to as being ~5 times that of the IC concentration.

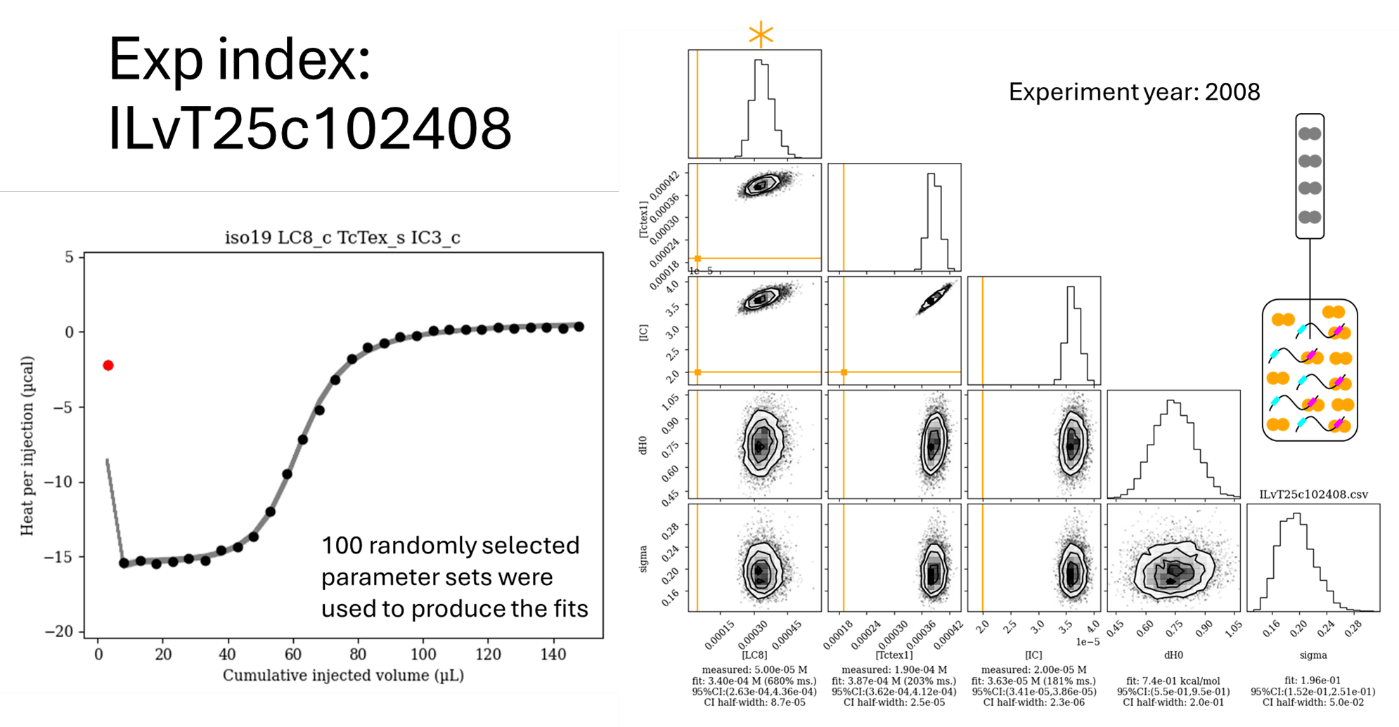

Figure S21. Isotherm ILvT25c102408 with the distributions of the fits of the concentrations and other nuisance parameters associated with the given isotherm. 100 random parameter sets from the posterior distribution were selected for drawing the fits on the isotherm. As shown in the cartoon, Tctex1 was titrated into a mixture of IC and LC8 for this experiment. The red points in the isotherm indicate injections that were disregarded in the fit. Orange bars on the corner plot correspond to values input as “measured”. The asterisk denotes that the LC8 concentration was not recorded for this experiment but was referred to as being ~5 times that of the IC concentration.

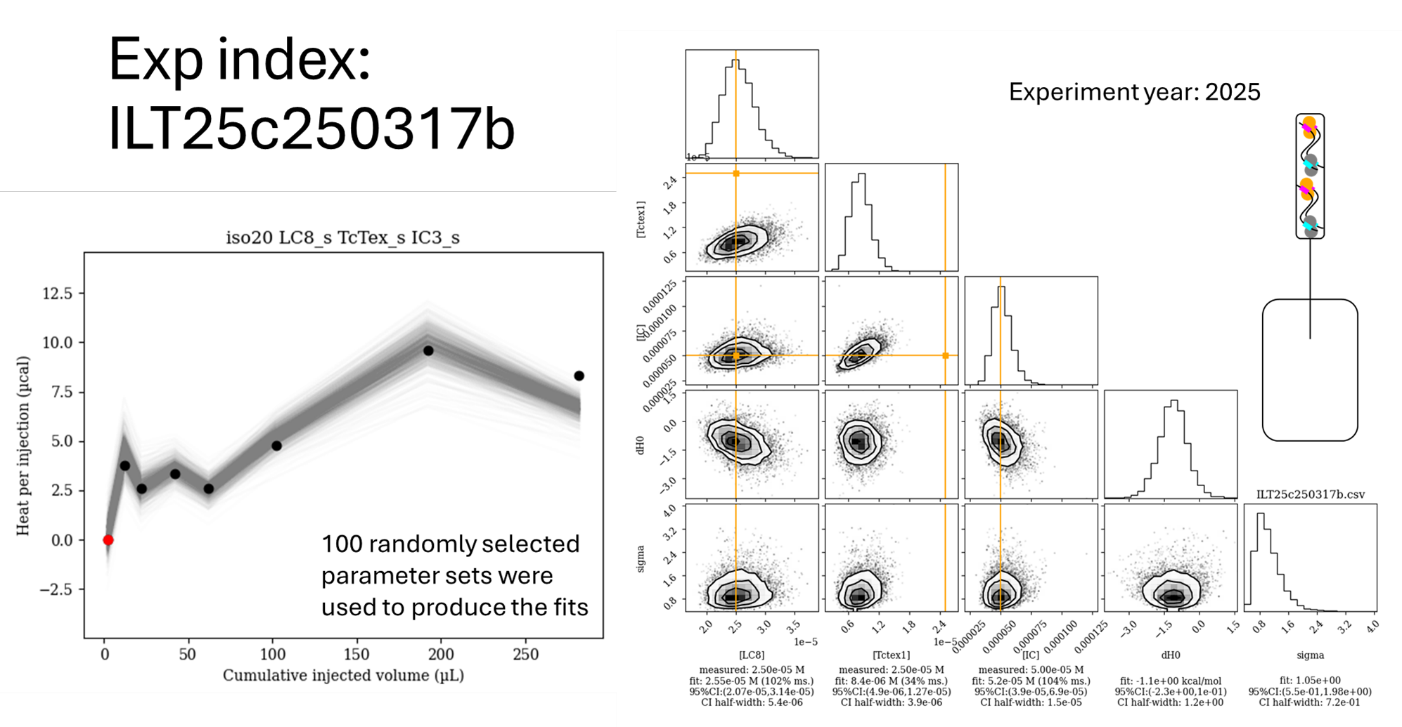

Figure S22. Isotherm ILT25c081408 with the distributions of the fits of the concentrations and other nuisance parameters associated with the given isotherm. 100 random parameter sets from the posterior distribution were selected for drawing the fits on the isotherm. As shown in the cartoon, a mixture of IC, LC8, and Tctex1 was titrated into buffer for this experiment. Note that the injection volumes vary across the run to continue measuring significant heats. The red points in the isotherm indicate injections that were disregarded in the fit. Orange bars on the corner plot correspond to values input as “measured”.

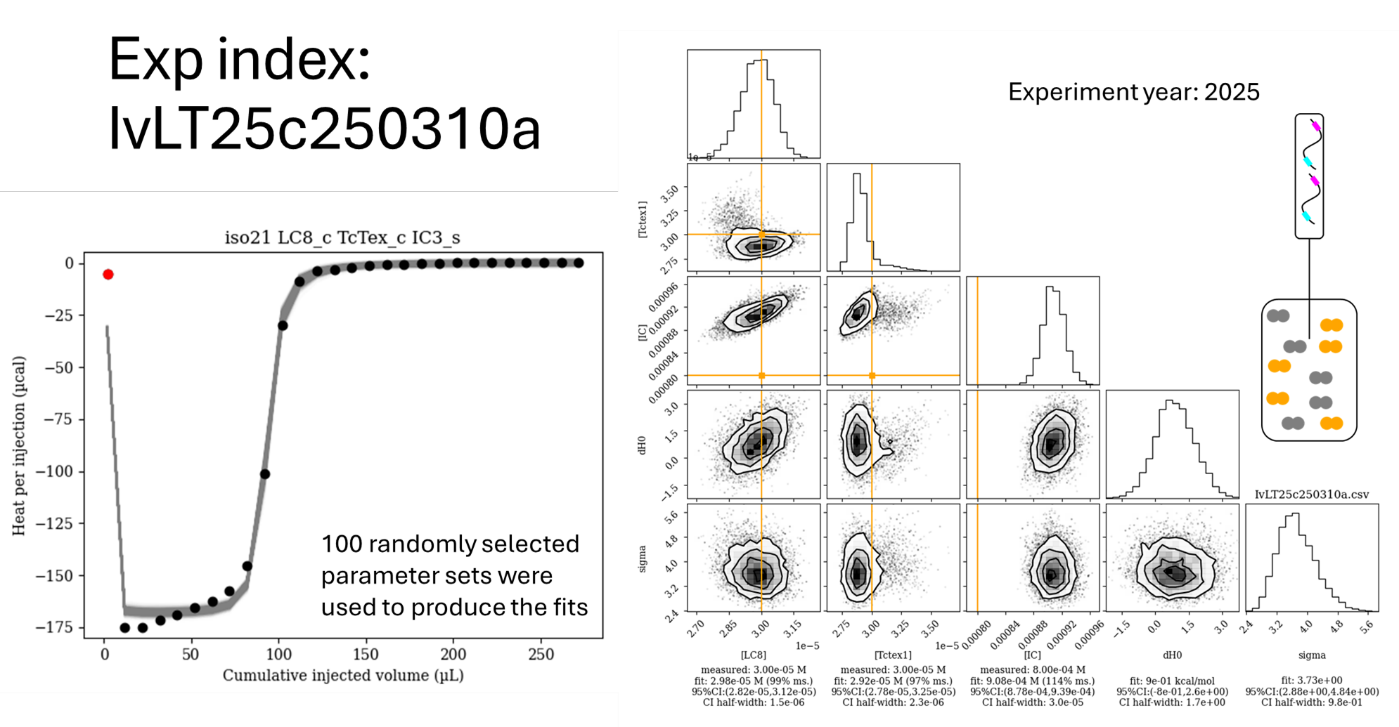

Figure S23. Isotherm IvLT25c250310a with the distributions of the fits of the concentrations and other nuisance parameters associated with the given isotherm. 100 random parameter sets from the posterior distribution were selected for drawing the fits on the isotherm. As shown in the cartoon, IC was titrated into a mixture of LC8 and Tctex1 for this experiment. The red points in the isotherm indicate injections that were disregarded in the fit. Orange bars on the corner plot correspond to values input as “measured”.

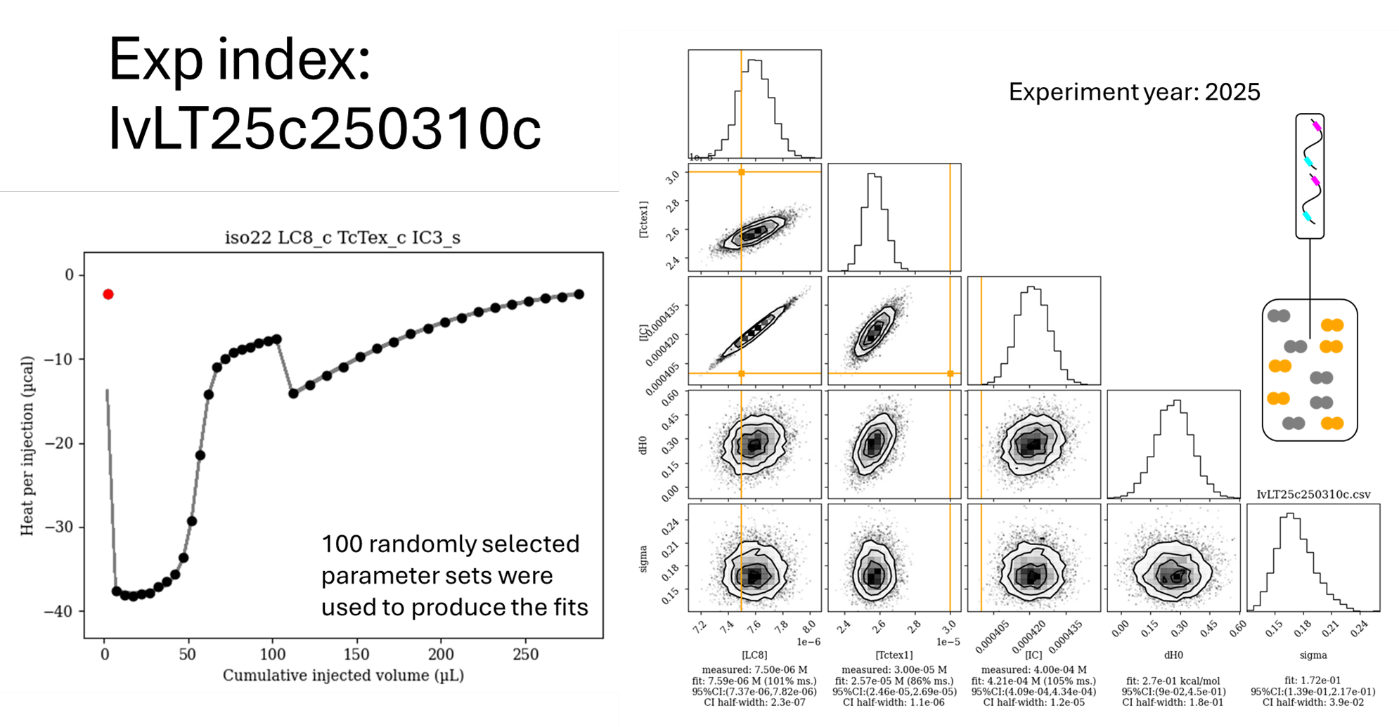

Figure S24. Isotherm IvLT25c250310c with the distributions of the fits of the concentrations and other nuisance parameters associated with the given isotherm. 100 random parameter sets from the posterior distribution were selected for drawing the fits on the isotherm. As shown in the cartoon, IC was titrated into a mixture of LC8 and Tctex1 for this experiment. Notedly, larger injection volumes were used for the second half of the experiment. The red points in the isotherm indicate injections that were disregarded in the fit. Orange bars on the corner plot correspond to values input as “measured”.

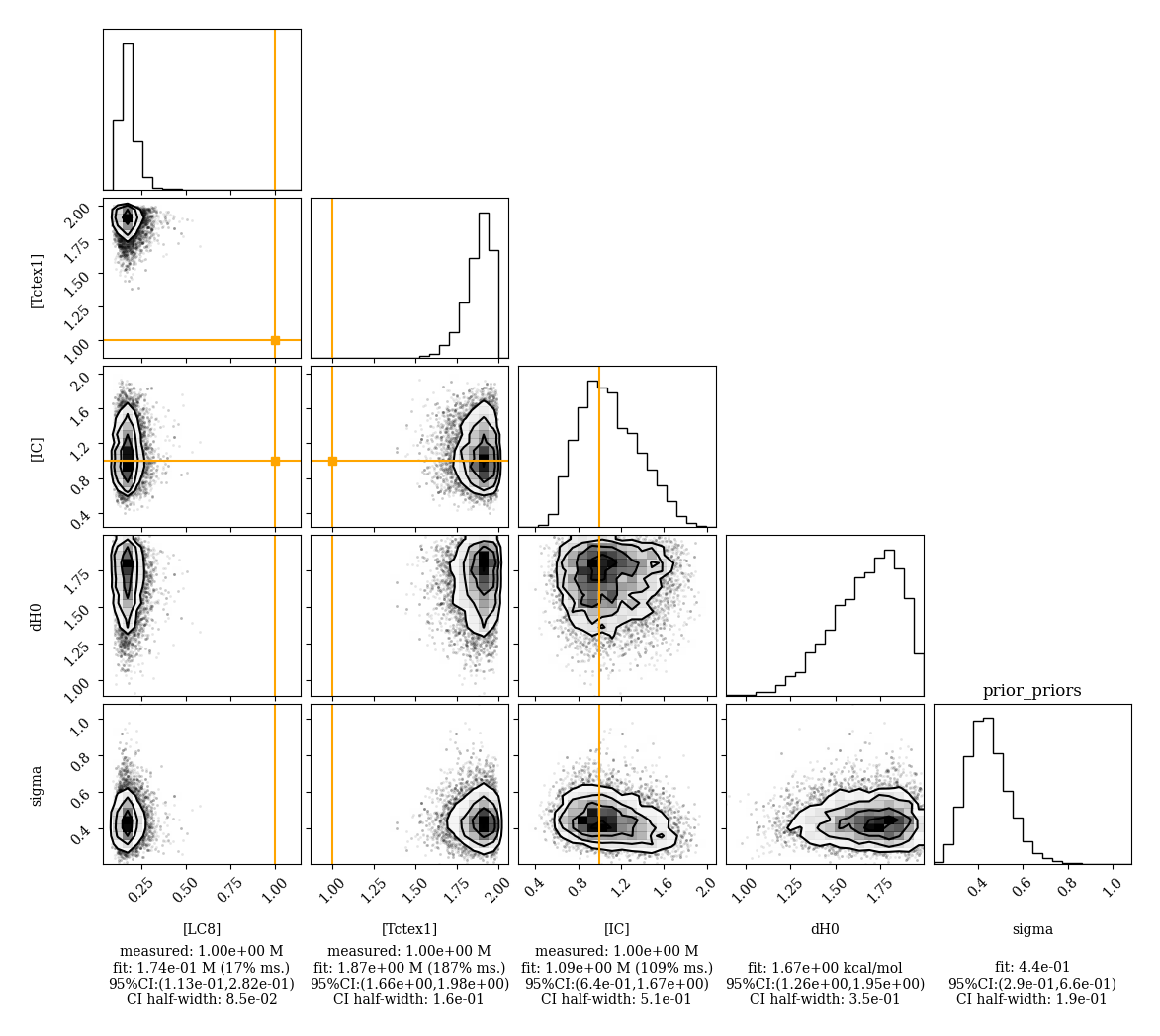
Figure S25. Distributions of the master prior which oversees the allowed prior ranges over which all of the nuisance parameters are allowed to vary. Orange bars correspond to 100% variation on a log-scale.
